# Rapid and Signal Crowdedness-Robust In-Situ Sequencing through Hybrid Block Coding

**DOI:** 10.1101/2022.11.16.516714

**Authors:** Tianyi Chang, Wuji Han, Mengcheng Jiang, Jizhou Li, Zhizhao Liao, Mingchuan Tang, Jianyun Zhang, Jie Shen, Zitian Chen, Peng Fei, Xianwen Ren, Yuhong Pang, Guanbo Wang, Jianbin Wang, Yanyi Huang

## Abstract

Spatial transcriptomics technology has revolutionized our understanding of cell types and tissue organization, opening new possibilities for researchers to explore transcript distributions at subcellular levels. However, existing methods have limitations in resolution, sensitivity, or speed. To overcome these challenges, we introduce SPRINTseq (Spatially Resolved and signal-diluted Next-generation Targeted sequencing), an innovative in situ sequencing strategy that combines hybrid block coding and molecular dilution strategies. Our method enables fast and sensitive high-resolution data acquisition, as demonstrated by recovering over 142 million transcripts using a 108 gene panel from 453,843 cells from four mouse brain coronal slices in less than two days. Using this advanced technology, we uncover the cellular and subcellular molecular architecture of Alzheimer’s disease, providing additional information into abnormal cellular behaviors and their subcellular mRNA distribution. This improved spatial transcriptomics technology holds great promise for exploring complex biological processes and disease mechanisms.

## Introduction

The well-ordered structural organization of cells and the intrinsic heterogeneity among cells are essential characteristics of multi-cellular organisms. Various spatial transcriptomic technologies have been developed to help understand the nature of such spatial information and its functional properties. In-situ mRNA capture-based approaches offer a spatially-resolved expression landscape (*1-5*). Nevertheless, these methods have trouble providing finer information in space, such as subcellular RNA localization, whereas imaging-based spatial transcriptomic approaches offer near-optical limit resolution (*6*). An ideal spatial transcriptomic technology needs to be sensitive, accurate, scalable, and robust. Such currently available approaches are divided into two major categories, fluorescent in-situ hybridization (FISH) (*7-11*) and in-situ sequencing (ISS) (*12-16*). Those prevalent approaches face major challenges due to time-consuming workflows, complex reagents, and complicated imaging setups. FISH-based approaches are typically built upon single molecule detection schemes that rely on high-power, large numeric aperture objective lens and on specialized imaging techniques to overcome a low signal-to-background ratio (SBR) problem, and that usually leads to a sacrifice in the size of imaging field of view (FoV). The length of the mRNA of interest is also limited since many probe binding sites are commonly needed. While ISS-based methods usually apply in-situ amplification to increase the SBR ratio, molecular crowdedness during amplification or optical imaging are major challenges. Another common issue of currently available methods, the long experimental time, greatly hinders the scalability needed to handle a high-quantity of or large-size samples.

Here, we introduce a new ISS-based technology that uses sequencing-by-synthesis (SBS) chemistry to speed up the cyclic reactions needed for highly efficient information acquisition. We created a hybrid block code that is both signal-crowdedness and error robust, and that has a high decoding efficiency. Physical dilution of signal is adopted to further counteract ISS’s intrinsic crowdedness issue. This method, SPatially Resolved and signal-diluted Next-generation Targeted sequencing (SPRINTseq), has greatly shortened the sequencing time it took (to within 9.5 hours) to profile a targeted transcriptome of a mouse brain coronal slice, and produced near optical diffraction-limit resolution. SPRINTseq produced a sub-micron precise, whole-slice-scale cellular atlas that contains subcellular location information for each transcript. Such an information-rich sub-cellular distribution of genes can be greatly affected by physiological conditions. Using a 108-gene panel, 4 slices of mouse coronal brain (from 2 normal mouse and 2 mouse with Alzheimer’s disease) from sample to data could be profiled in 2 days, covering 453,843 cells and 142,957,485 transcripts. We found that the degree of subcellular mRNA dispersion increased as glia cells activated in Alzheimer’s disease. We also found that the mRNA distribution produced changes in orientation within the amyloid microenvironment. Additionally, the high heterogeneity we found within inhibitory neurons may correlate with cell-type-specific responses during disease.

## Result

### Principle, operation, and performance of SPRINTseq

A SPRINTseq experiment consists of two major parts, in situ barcoded clonal amplification and ISS. We constructed an integrated setup to automatically control fluidics, temperature, mechanical motion, and fluorescence imaging (Fig. 1A, Fig. S1 & S2). A slice of tissue was placed in a microfluidic flow-cell and various reagents were programmed to flow through it. A set of padlock probes we designed directly hybridized mRNAs with high specificity and then ligated to form a circular DNA template for rolling circle amplification (Fig. 1B). In about 10 hours, targeted genes were converted in situ into individual nanoball clones of corresponding barcodes. Each padlock probe contained two identical barcode blocks to double the signal of the sequencing reaction (Fig. S3), which came from fluorescent-labeled, reversible terminator nucleotide substrates. Each barcode block consisted of a gene-specific barcode and a universal sequencing primer binding sequence.

**Fig. 1.**
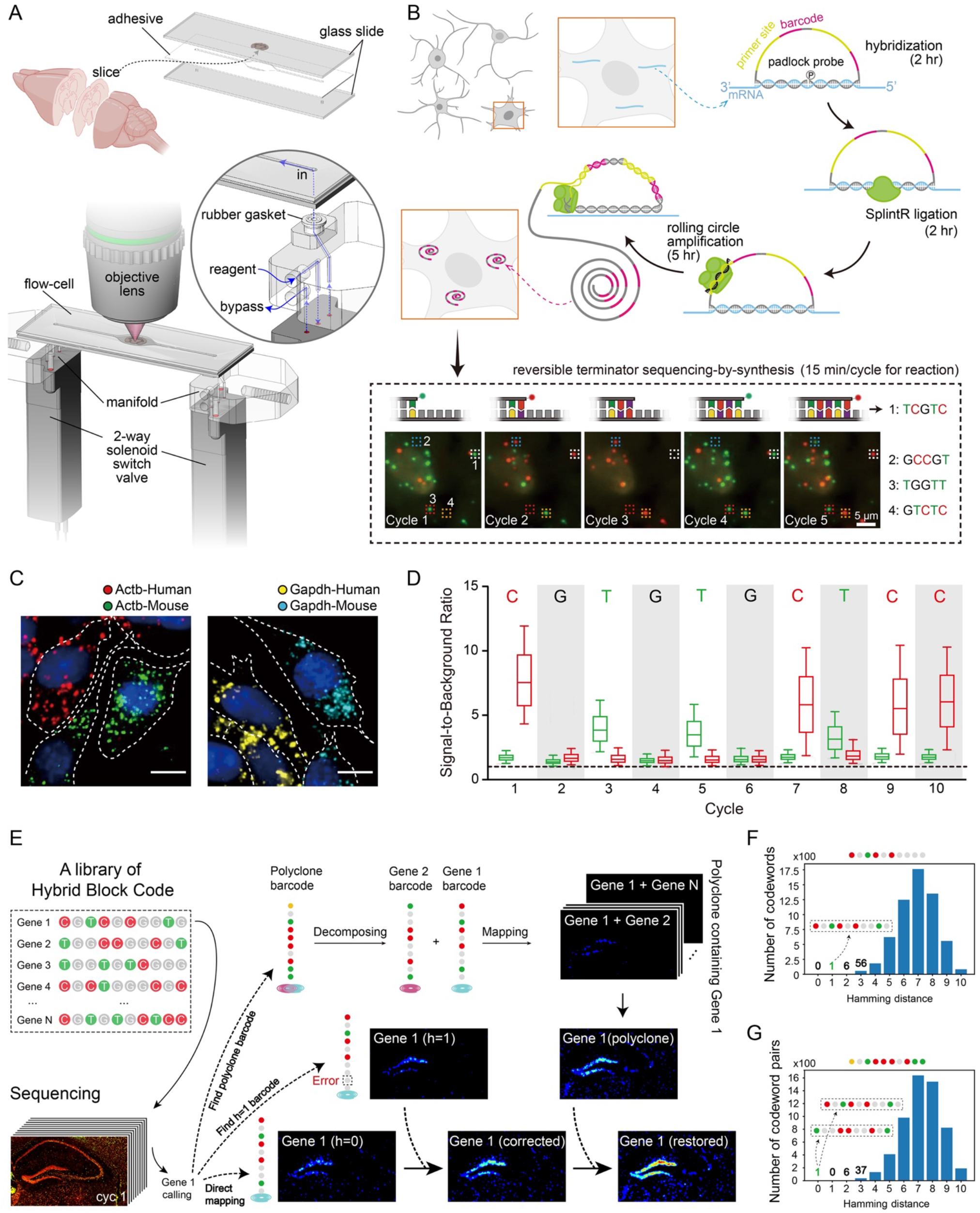
Workflow and performance of SPRINTseq: in-situ sequencing from sample to data within 20 hours. **(A)** In-situ sequencing automation. Each mouse brain tissue slice was mounted on a slide and each slide was assembled into a flow cell and mounted between 2 manifolds. The manifolds were controlled by 2-way solenoid switch valves that controlled the fluidic routes. The reagent flowed directly into the flow cell while a bypass prevented reagent cross-contamination. **(B)** Schematic workflow: in situ barcoded clonal amplification and in situ sequencing. Probe hybridization, ligation, and rolling circle amplification were performed sequentially and took about 10 hours total. Each probe contained two identical “sequencing primer + barcode” blocks to increase the signal-to-background ratio. Barcodes were sequenced in situ using 2-color, reversible terminator sequencing-by-synthesis chemistry. Scale bar: 5 μm. **(C)** SPRINTseq specificity demonstrated by inter-species gene detection on a human (HEK293T) and mouse cell (3T3) co-culture system. Scale bar: 10 μm. **(D)** Characterization of 10-bp barcode sequencing. No obvious dephasing or signal decay was observed throughout 10 cycles. The boxes show the interquartile range, the lines in the boxes are the medians, and the bars show min/max values. The horizontal dashed line is SBR = 1. **(E)** Hybrid block codes achieve error-correctable and crowdedness-robust encoding. The barcodes are highly orthogonal between each other. A specific gene is mapped by looking for its exact barcode sequence and the one with correctable error (e.g., Hamming distance [h] = 1). Also, all composed barcodes (composite codewords) containing that gene are corrected for mapping. By counteracting signal overlapping and sequencing error, a significant number of signals are rescued by this encoding strategy. **(F)** Error correction. When a sample readout sequence “CGTCGCGGGG” (one base error) is aligned with all barcodes (including an original barcode and a composed barcode) in the library, only one barcode (CGTCGCGGTG, Gene 1) with a Hamming distance equal to 1 is found, thus this sequence was mapped to Gene 1 and the error was corrected. **(G)** Polyclone barcode decomposition. When a sample readout sequence “AGTCCCGCTT” (a composed barcode containing two barcodes) is aligned with all the composed barcodes in the library, only one barcode pair (Gene 1: CGTCGCGGTG and Gene 2: TGGCCGGCGT) with a Hamming distance less than or equal to 1 exists. Thus, this polyclone (overlapping amplicons) can be decomposed and mapped to Gene 1 and Gene 2.

Because of its fast cyclic reaction time (Fig. S2; 15 min/cycle includes tissue blocking, substrate incorporation, and fluorophore cleavage steps), we used mature 2-color reversible terminator SBS chemistry for barcode readout (Fig. 1B). With an approximate 2.2 × 10^7^ pixel/s imaging speed, the sequencing process took only about 9.5 hours for a 10-base barcode reading on a coronal slice of mouse whole brain. The images were further processed for base calling and cell segmentation (Fig. S5 & S6). Common imaging background noise caused by non-specific binding of the fluorescent nucleotide substrate was clearly reduced when free thiol groups in the tissue were blocked (Fig. S3A,B). Also, with multiple-barcode-block design (Fig. S3C-F), the SBR of SPRINTseq was significantly greater than those FISH-based methods (Fig. S7). Altogether, signal with sufficient SBR can be generated even on high auto-fluorescence tissue samples (Fig. S4). Amplified barcode clones were stably attached on the sample and drifted negligibly during the whole experimental process (Methods). Using only one probe per gene, targeting sensitivity was as high as 38% (Fig. S7D-G), mainly due to a low loss of amplified clones and no reverse transcription step in the protocol. In addition, during cyclic sequencing reactions, SPRINTseq exhibited high specificity (Fig. 1C) with very low signal decay or dephasing (Fig. 1D, Fig. S8).

We designed a highly efficient and orthogonal coding scheme, ‘hybrid block code’, that encodes barcodes in a way that tolerates signal crowdedness and corrects sequencing errors (Fig. 1E). We encoded 2^k^ genes using n-bit (n > k) codewords created from 0/1 string-based block code, and used the redundant bits to correct errors (Fig. S9). Moreover, we folded n-bit binary codewords to (n/2)-bit double-binary codewords by combining every two bits into a dual channel bit, thus shortening signal acquisition time. The dual-channel “1” signal at the same bit was avoided to reduce the overall signal density and to improve orthogonality between codewords. We then selected a subset of all codewords, among which almost any two could be uniquely split to enable the decomposition of overlapping signals. Hybrid block code naturally fits 2-color reversible terminator SBS. To convert the codeword into a DNA barcode sequence, we used the bases ‘C’ and ‘T’ to represent “1” in the two channels and base ‘G’ to represent “0”. Base ‘A’ was not used in the barcodes. After sequencing, all barcode reads were aligned against a library consisting of the original barcodes and all composed barcodes (e.g., CGG + TCG = ACG). When aligning a barcode that had one sequencing error against the library, only one barcode with a Hamming distance equal to 1 was obtained, thus achieving error correction (Fig.1 F). Polyclone (overlapping amplicons) barcodes were aligned to all possible polyclone barcodes (all barcode pairs) in the library. Thus, a unique polyclone barcode with a Hamming distance equal to or less than 1 was found, and that allowed us to decompose the signal from overlapping amplicons (Fig.1 G). Using barcodes called from an image sequence built with that design resulted in a significantly higher average Shannon entropy per image than those of existing methods (Table S3). Our refined experimental process and efficient coding scheme enables SPRINTseq to profile a coronal slice of mouse whole brain within 20 hours and with high sensitivity and accuracy.

### Relieving crowdedness issues yields high quality ISS

Signal crowdedness, a major challenge for ISS, occurs when numerous amplified signal spots fill the limited space within a cell and thus limit the total read count. In a swift scanning scheme, the locations of amplified clones after 2D-stacking projection are prone to signal overlap in crowded regions. In hybrid block code, though, signals are diluted by using the ‘bit-of-silence’ and ‘optical dilution’ is realized. Only a fraction of the barcodes was fluorescently labeled in each sequencing cycle (Fig. S9C). Such signal sparseness can be finetuned by the number and position of the ‘0’ signal in the barcodes, thus staggering the signals from highly expressed genes in different cycles.

However, the optical dilution ratio has an upper limit within a certain number of sequencing cycles as the proportion of silent channels is limited by the Hamming weights of codewords. Inevitably, signal co-occurrence between highly-expressed genes will happen as more genes that need to be identified are included in the panel (Fig. 2A). In some cases, a signal from one super-highly expressed gene is so dense that the spots form ‘plaques’ and the signals from other genes are covered and their information is unextractable. This crowdedness is caused by too many amplicons from highly abundant genes. Also, some amplification events may be precluded in such a physically crowded environment, thus causing an even greater bias in gene profiling. Optical dilution does not alleviate this issue.

**Fig. 2.**
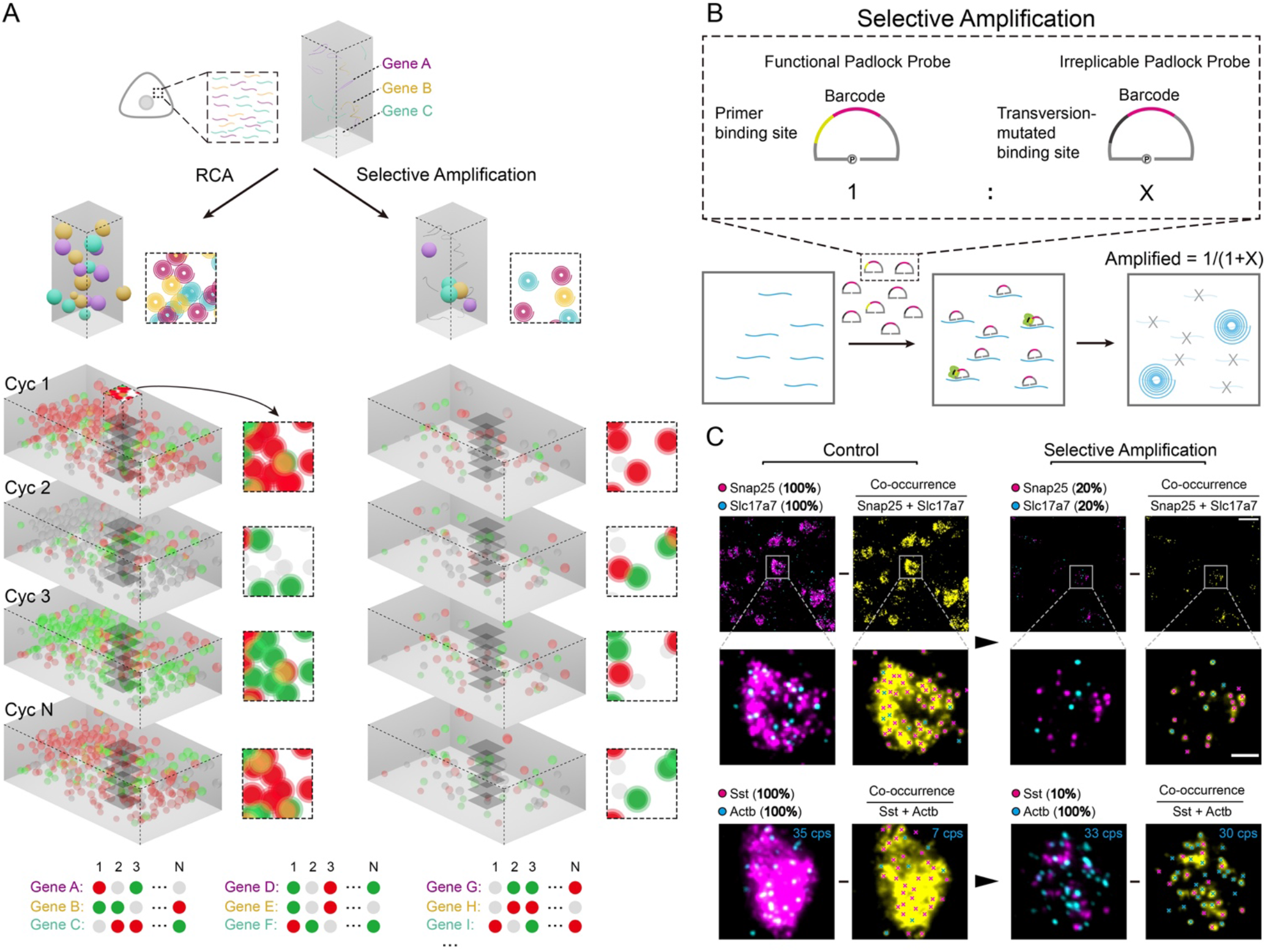
SPRINTseq uses signal dilution to relieve signal crowdedness. **(A)** Spatial crowdedness and physical dilution of signals. As signals from rolling circle amplified (RCA) clones were projected onto a 2-D plane during focal-stacking, the signals inevitably overlapped in crowded regions and became unrecognizable, thus causing information dropout. Through selective amplification, signals from partial transcripts were physically eliminated, thus leaving a sparser environment. (**B)** Selective amplification. The proportion of amplification of specific highly expressed genes can be controlled by mixing normal padlock probes and sequencing-primer binding-site transversion probes (irreplicable), which achieve selective gene masking during amplification and yields a desired ratio. **(C)** Signal rescue through selective amplification. Respective signals from Snap25/Sst (magenta) and Slc17a7/Actb (cyan) can be obtained through another sequencing cycle. Because at least one base of the respective gene barcodes is designed to be the same (cycle and channel),a crowdedness situation (co-occurrence, yellow) is simulated. Percentages represent the remaining fractions of gene signals after selective amplification. The crosses represent the recognizable signals based on local maxima identification. In the example image, 20% (7/35) of Actb reads can be extracted during co-occurence with all Sst amplicons, while 91% (30/33) of the reads can be extracted during co-occurrence after masking 90% of the Sst amplicons. Scale bars: 20 μm for the base images and 5 μm for the insets.

We used a selective amplification strategy to relieve physical crowdedness. For those highly expressed transcripts, we doped the padlock probes with irreplicable ones to uniformly dilute their clonal populations. Thus, partial transcripts of such genes are physically eliminated and other genes are able to be better amplified and read (Fig. 2B). To demonstrate selective amplification, we simulated an inevitable situation in which multiple genes are tested in a panel: two genes with dense signals appear in the same reaction cycle and channel. Snap25 and Slc17a7 are highly expressed genes in excitatory neurons and when their signals appeared together, their clonal puncta joined together to form plaques in the images and the puncta were obscured (Fig. 2C). After masking 80% of both genes, each of their transcript clones were clearly differentiated. Though the masked signals’ locations could not be recovered, the quantities of transcripts could be re-calibrated by multiplying the dilution ratios in gene-by-count expression matrix (Fig. S10). In another example, Sst, a GABAergic neuron subtype marker gene, dominated the expression in that cell type. The puncta of clonal signals from other genes such as Actb were difficult to discern. After diluting the Sst signal 10-fold, it was possible to digitally count mRNA clones, and many signals, including those of Actb and Sst, were rescued. Only 20% (7/35) of the Actb reads could be extracted when the Actb signal co-appeared with Sst’s. After masking 90% of the Sst amplicon, more than 90% (30/33) of the Actb reads were extracted. Given prior knowledge, we can adjust the physical dilution ratios for genes with different abundances. This ratio needs to be high enough to identify clonal signals in highly expressed cells, but not so high as to cause false-zeroes in lowly expressed cells. Optical dilution and selective amplification are functionally complementary. The final dilution fold is the product of the physical dilution fold from selective amplification and the optical dilution fold.

For regions that are still crowded after two-level dilution, the polyclonal signals that result mainly from the co-localized 2-D projections of two amplicons, can be decomposed into separate barcodes in hybrid block coding and their signals can be rescued (Fig. 1E,G). By combining optical dilution, physical dilution, and polyclone decomposition, SPRINTseq effectively relieves the signal crowdedness issue in ISS.

### Single cell profiling with subcellular resolution of mouse brain coronal slices

We profiled mouse brain coronal slices (approximately 10.2 × 7.6 mm^2^) at subcellular resolution using a 108-gene panel (Fig. 3A). The genes included both marker and disease-related genes selected for cell classification and status characterization (Extra Table 1) (*17*). To encode these genes, we designed a 10-base barcode set based on the hybrid block code (Fig. S5A-C). The minimum Hamming distance between any barcode pair in the 108-gene panel was 3, and the minimal Hamming distance between almost any (> 99.8%) barcode pairs that containing polyclone barcode was 3, which ensured error correction and polyclone decomposition. The silent bit (G) content of 52.4%, achieved a 23.8% optical dilution ratio while keeping a sufficient Hamming weight in codewords. Furthermore, selective amplification applied to seven highly-expressed genes (Actb, Mbp, Cst3, Penk, Snap25, Slc17a7, and Sst) to further reduced the clone number by 54.4%. As a result, about 10% of all signals were acquired per sequencing cycle, significantly diluting crowdedness. While 49.8% of all monoclonal reads were rescued from error correction, 31.6% of the total reads were rescued from polyclone decomposition.

**Fig. 3.**
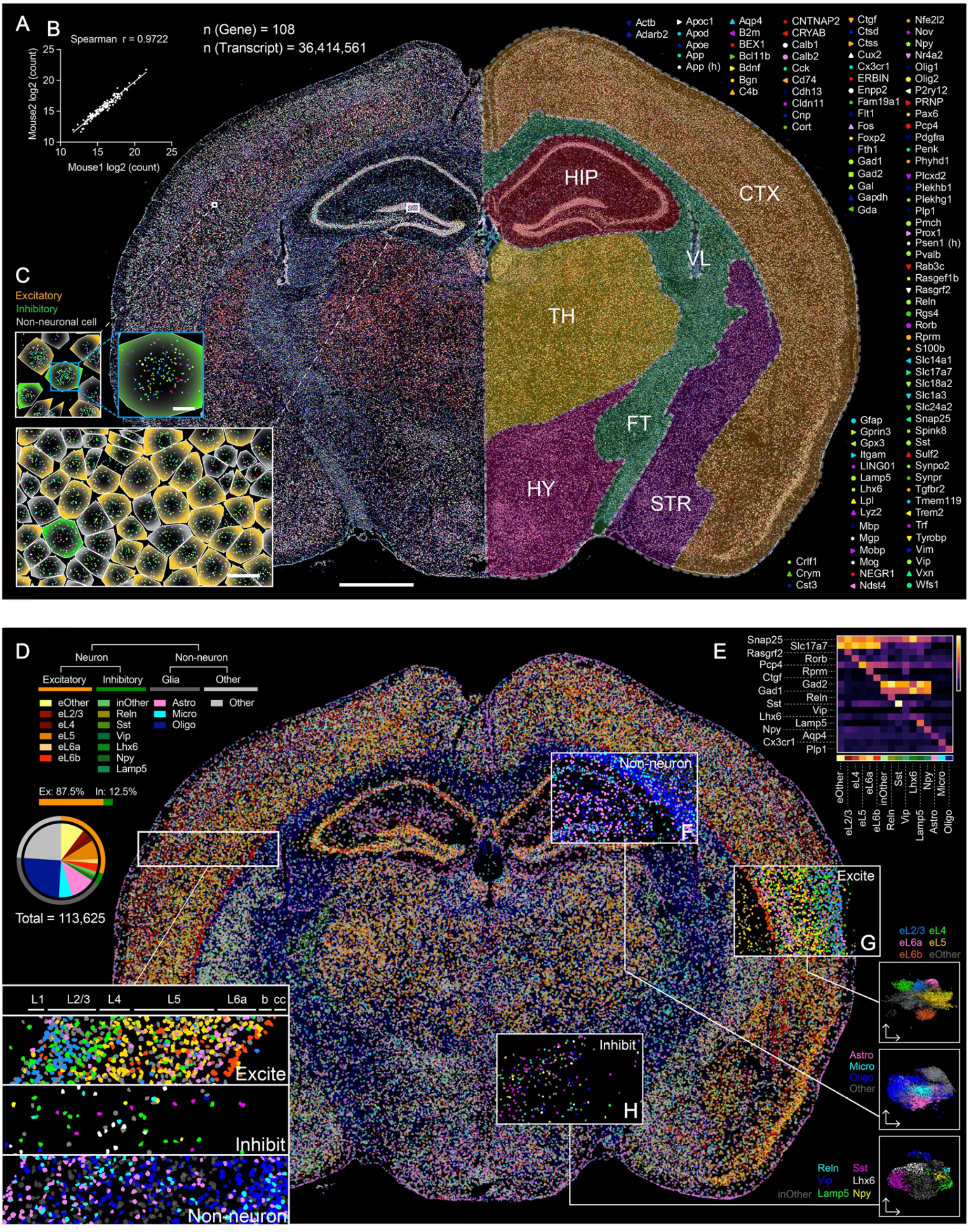
Single cell profiling with subcellular resolution of mouse brain coronal slices. **(A)** The spatial landscape of 108 genes called by 10-cycle sequencing in mouse brain. Brain regions shown in the right half, the brain was defined by several regional marker genes (Slc17a7, Gad2, Fth1, Enpp2, Pcp4, Pmch), CTX: cortex, HIP: hippocampus, VL: lateral ventricle, TH: thalamus, FT: fiber tract, HY: hypothalamus, STR: striatum. Scale bar: 1 mm. **(B)** Spearman correlation of two mouse sequencing replicates, the brain slices were selected at the same position and the gene panel was the same. **(C)** Enlarged images showing gene locations (dots color/shape-coded to the genes) and cells from the insets in **(A)**. Cell border was showed by lines with different colors: orange for excitatory neuron, green for inhibitory neuron and gray for non-neuronal cell. Scale bar: 2 μm for the single cell image and 10 μm for the multiple cell images. **(D)** Cell composition and spatial projection of a brain slice. Marker gene expression levels and Louvain shared nearest neighbor clustering identified 17 cell types (16 defined cell types and 1 other cell type shown in upper left corner), and their proportions are shown in the pie chart. Each color represents one unique cell type. The enlarged images at the bottom left show projections of all 17 cell types at the same locations in the cortex. The anatomical structure is labeled on top beginning with L1 (L: layer in the cortex, cc: corpus callosum) **(E)** Marker gene expression heatmap by each cell type. The cell type color code is the same as in **(D). (F-H)** Spatial projections of non-neuronal cells (astrocytes, microglia, and oligodendrocytes), excitatory neurons (eL2/3, eL4, eL5, eL6a, eL6b and others), and inhibitory neurons (Reln, Sst, Vip, Lhx6, Lamp5, Npy, and others), as well as their cell clustering visualizations through uniform manifold approximation plots.

We identified 16,606,784 raw reads from the brain slice, and that number increased to 36,414,561 after selective amplification was re-calibrated. Probe binding false positives and gene mapping were 0.05 and 0.02 events per cell, respectively. The border of each general brain region was drawn using corresponding marker genes (Fig. 3A). Highly accurate and specific expression patterns were consistent with the in-situ hybridization results in the Allen Brain Atlas (Extra Supplement PDF). Brain replicates also showed high concordance at both whole and regional brain levels (Fig. 3B, Fig. S11).

We determined cell nuclei centroids using DAPI staining, calculated cellular segmentation with nucleic images, and assigned sequenced RNA reads to their nearest nucleus centroid (Fig. 3C, Fig. S6). We obtained 146,137 cells in total and 113,625 (77.8%) passed quality-control. According to the major marker genes, cells were categorized into three major populations: excitatory neurons (Slc17a7+), inhibitory neurons (Gad1/Gad2+), and non-neuronal cells. Each population was then subdivided into detailed clusters through Louvain shared nearest neighbor clustering and the resulting 17 types were spatially projected to their original positions (Fig. 3 D & E, Fig. S12-13). Non-neuronal cells were divided into astrocyte, microglia, oligodendrocyte, and other cell types (Fig. 3F). Astrocytes and microglia are scattered across the whole brain, but astrocytes are more frequently distributed at tissue edges and the hippocampal region. Oligodendrocytes have a high density in the fiber tract region. Neuronal cells were classified according to their positions in the cortex (for excitatory neurons, Fig. 3G) and their subtypes (for inhibitory neurons, Fig. 3H). Excitatory neurons in the cortex can be clearly divided into six layers. They are also widely distributed in the hippocampus and thalamus. Inhibitory neurons are sparsely distributed in the cortex, striatum, hippocampus, hypothalamus, and reticular nucleus region of the thalamus.

SPRINTseq offers an informative subcellular distribution and location of genes, as well as their correlations to other spatially distributed components. We assessed the degree of the mRNAs’ subcellular dispersity using the average distances to their centroids, and also calculated the mRNA average distance to the nucleus centroid (Fig. S14). We then used those two parameters as coordinates for each gene in the panel and classified them all into one of three quadrants.

The mRNAs of the genes in the first quadrant were widely dispersed within the cell and far from the nucleus. From that we inferred that they were diffused throughout the cytoplasm and expressed the protein needed inside the cell. Actb is one typical example of this type. The mRNAs of the genes in the third quadrant, such as Ctgf, were likely to be distributed on the endoplasmic reticulum near the nucleus and to express membrane proteins or secreted protein. Genes appearing in the fourth quadrant showed a certain polarity in their mRNA distributions; genes including Snap25 and Slc17a7 were in this category. The mRNA distribution polarity of signal-transduction-associated genes may be related to the polarity of neurons and other cells in the brain. No genes appeared in the second quadrant.

### Spatial profiling of the mouse brain with Alzheimer’s disease

The 108-gene panel included many genes associated with Alzheimer’s disease (AD), which is also pathologically associated with the distinct spatial distribution of amyloid plaques (Amyloid β) that accumulate in the cortex and hippocampus, especially during aging (*18*). We used SPRINTseq to examine brain coronal slices from 10-month-old APP/PS1 and normal mice. First, the genes’ overall expression levels were similar between AD and normal mice at the whole brain slice and regional levels (Fig. S15A). Next, we investigated gene-expression changes around amyloid plaques, whose typical size was over 100 μm in diameter. We aligned the Amyloid β immunofluorescence images to our SPRINTseq images and defined the plaques and their surrounding areas as ‘pan-plaque regions’ (Fig. 4A). Spatial correlation analysis between all genes and Amyloid β in those regions showed that many genes in our panel were potentially responsive to Amyloid β (Fig. 4B). The most relative genes included Gfap, Tyrobp, Ctsd, B2m, and Apoe, which express mainly in reactive astrocytes and microglia cells (*19, 20*). The aggregation of those genes was confirmed by raw sequencing images (Fig. 4B), and the ranking of spatially-correlated genes were independent of the bin size selected for analysis (Fig. S15B).

**Fig. 4.**
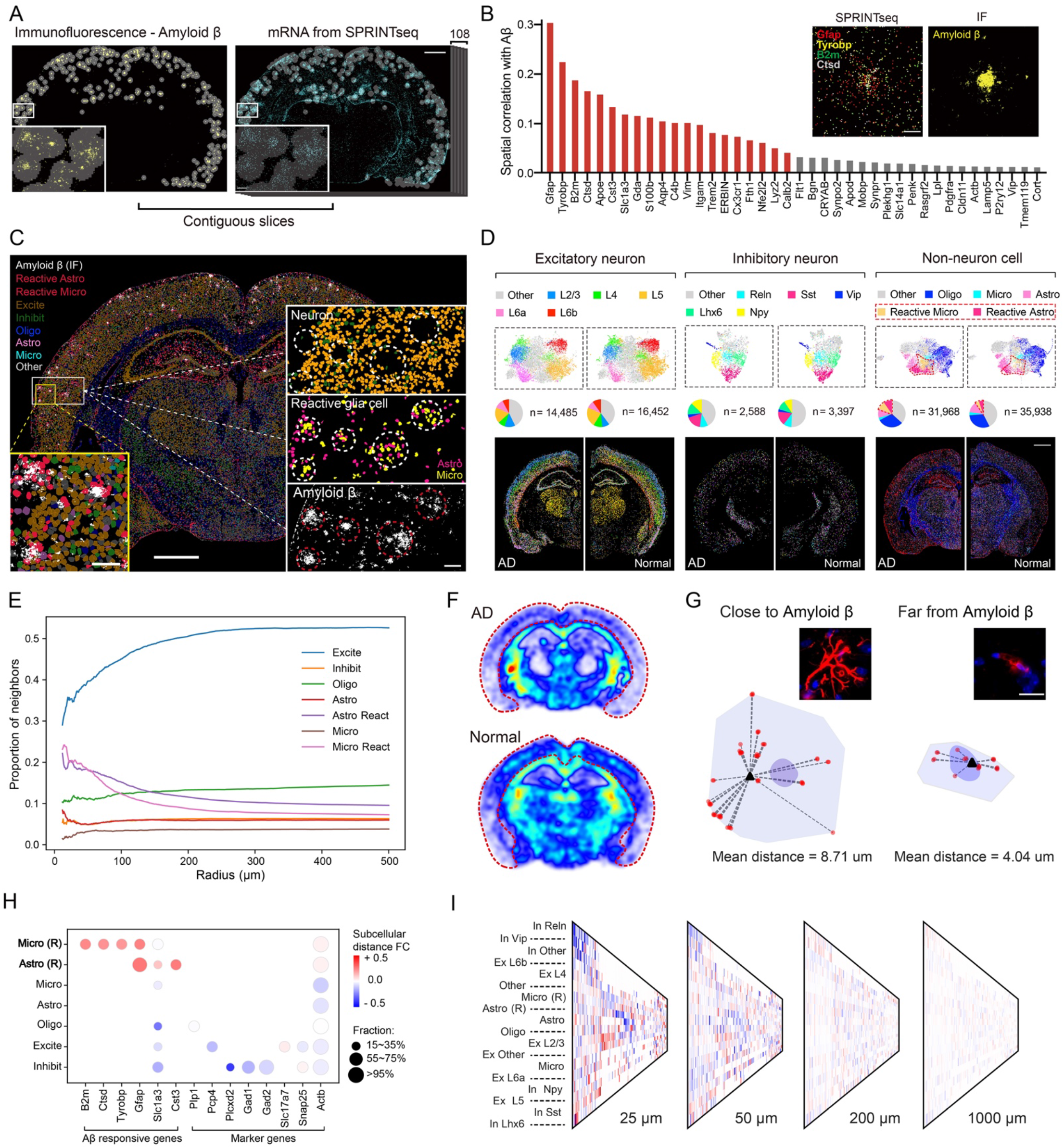
Spatially-resolved cellular and subcellular changes of mouse brain with Alzheimer’s disease (AD). **(A)** Immunostained amyloid plaque (amyloid β) and SPRINTseq analysis on contiguous slices. After alignment, the amyloid plaque positions were localized on the sequencing image. The ‘Pan-plaque region’ (regions circled in white) was defined around amyloid, and spatial correlation was calculated between all 108 genes and the amyloid plaque, respectively, in the pan-plaque regions. Bin size: 50 × 50 μm. Scale bars: 80 μm for enlarged images and 1 mm for brain slice images. **(B)** The most enriched genes, as shown by spatial correlation analysis. The top 20 genes in the graph are shown by red bars. The upper right images show the top 4 genes’ (Gfap, Tyrobp, B2m, and Ctsd) actual aggregations (left, sequencing result) and at the same positions with amyloid plaque (right, immunofluorescence [IF], contiguous slice after alignment). Scale bar: 40 μm. **(C)** Whole brain spatial projection for general cell types. Amyloid plaque IF in a contiguous slice is shown along with excitatory and inhibitory neurons and non-neuronal cells: oligodendrocyte (Oligo), astrocyte (Astro), microglia (Micro), and other. Reactive glia cells (responding to amyloid plaque) are labeled in red (lower left image). The 3 enlarged images on the right show neuron and reactive glia cells and the amyloid plaque distributions at the same position. Scale bar: 1 mm for the whole brain image, 100 μm for lower left image and the multiple cell images on the right. **(D)** Normal and AD mouse brain direct cell composition comparison using combined classifications of the 16 cell types (see Fig. 3F-H). Scale bar: 1 mm **(E)** Relative cell density change as a function of distance to amyloid plaque. The measurements start at 11 μm. **(F)** Oligodendrocyte density comparison of AD and normal mouse brain. The cortex region is encircled by a red dotted line. Oligodendrocyte density decreased globally in the AD mouse brain. **(G)** Diagram of subcellular dispersion. The photos show reactive astrocytes with Gfap protein in red and DAPI in blue. IF results confirm that subcellular dispersion of reactive astrocytes differed based on their distances from plaque. In the corresponding diagrams, the red dots represent Gfap mRNA, the black triangles are Gfap mRNA centroids, the pale purple shows cell body, and the dark purple shows cell nucleus. Scale bar: 10 μm. **(H)** Heatmap of the degree of subcellular dispersion change. Y axis shows different general cell types as listed in (C). The values are the subcellular distance of cells close to amyloid plaque divided by that of cells far from amyloid plaque. (R): reactive, FC: fold change. **(I)** The orientation pattern as a function of distance. Y axis shows different cell types. The orientation of mRNAs was profiled in the cells whose distance to the nearest Amyloid β was 25 μm, 50 μm, 200 μm and 1000 μm, respectively.

At the cellular level, our SPRINTseq results showed that neurons were depleted from the Amyloid β while the reactive glia cells, including microglia and astrocytes, were aggregated around it (Fig. 4C). To directly compare the cell compositions of AD and normal brains, we combined both data sets and classified them together before projection (Fig. 4D, Fig. S15C). The proportion of reactive glia cells in the AD mouse brain was more than twice that in the normal mouse brain. For other cell types, although the difference in quantity between AD and normal brains was marginal, the local densities showed different spatial dependencies to Amyloid β and those changes were not random (Fig. 4E, Fig. S15D). Clearly, the reactive glia cells were clustered favorably around Amyloid β, within 200 μm, which reflected their roles of responding to microenvironmental changes and plaque clearance. The reduction of neuronal density near Amyloid β reflected neuron apoptosis in a plaque-rich microenvironment. Notably, the pathological effect of AD is not only associated with the local microenvironment but can be scaled to larger regions. For example, oligodendrocytes did not show significant density changes around Amyloid β, but their presence in the cortex region was generally less in the AD than in the normal mouse brain (Fig. 4F, Fig. S15E).

SPRINTseq extended the analysis’ spatial resolution to the subcellular level, enabling us to find many cells that showed distinct differences that were related to the cellular distance to Amyloid β. In reactive glia cells, the subcellular dispersion of mRNAs was increased in Amyloid β adjacent cells (distance threshold: 25 μm) than in more distant cells (Fig. 4G & H). This was largely because the cytoplasmic size of reactive glia cells near Amyloid β increased because of increased and elongated surface bumps. Disease-associated microglia (DAM) have been reported to play a vital role in AD and the function is conserved in mice and human (*21, 22*). Interestingly, the Amyloid β adjacent reactive microglia we found may largely correspond to reported plaque-phagocytic microglia (XO4+) in gene expression pattern (Tyrobp+, Apoe+, B2m+ and low Cx3cr1) and behavior (spatially enriched around plaques) (*23*). The enlarged cell body might suggest their function in phagocytosis and other regulation process within the micro-environment, whereas XO4-microglia can’t migrate towards plaques and thus are far away from plaques. The spatial information can be used as a new dimension for confirming plaque-phagocytic microglia.

Additionally, in a cell that is directly adjacent to a plaque, the location of one of its gene’s mRNA orientations with respect to the nearest Amyloid β can characterize the relative proximity of that gene. So, we calculated the included angle between the mRNA’s orientation, the nucleus centroid, and Amyloid β’s direction. A smaller angle indicated that the mRNA tended to be closer to its nearest plaque (Fig. S16A). Changes in the distance to the nucleus can also be calculated and combined with the orientation calculation to better describe the mRNA’s tendency to approach or retreat (Fig. S16B). Much heterogeneity was found within cells especially neurons, suggesting a different cell response pattern during disease (Fig. S16C). Such orientation was dependent on the distance between a cell and amyloid (Fig. 4I, Fig. S16D). This orientation feature pattern disappeared at longer distances, as the effect of Amyloid β on distant cells became weaker. Altogether, Amyloid β might generally affect its nearby cell morphology and potentially alter sub-compartment architecture in various cell types.

## Discussion

SPRINTseq is an intrinsically high-speed spatial sequencing method that can finish a mouse brain coronal slice profile at subcellular resolution within 20 hours, including both sample preparation (10 hours, and several slices can be prepared in parallel) and cyclic sequencing (9.5 hours). Due to a combination of highly effective barcode coding and robust SBS chemistry, SPRINTseq is significantly faster than most other approaches. Besides, we developed three orthogonal approaches to relieve the conventional ISS’s signal crowdedness issue. Among those approaches, optical dilution holds great potential and deserves further examination. For example, with constant codeword Hamming weights, more silent bits can be gained by increasing the barcode length. Multiple padlocks with different barcodes can also be applied to detect more transcripts of highly expressed genes, if necessary, which resembles selective amplification but is lossless in spatial information.

One major advantage of this method is the single-molecule high spatial resolution that gives sub-cellular localization information. This is important for addressing questions of cellular interactions in the context of gradients or proximity to specific tissue features such as disease lesions. In mouse models of Alzheimer’s disease, subpopulations of microglia with distinct transcriptomic phenotypes have been identified via single-cell RNA-sequencing (*24*). These seemingly AD-associated microglia bearing stronger inflammatory signatures and were hypothesized to engulf Amyloid β plaques (*24*). We described the distribution of different types of cells near Amyloid β in Alzheimer’s disease mouse model, and we identified reactive glia cells with a distinct expression pattern that are responsive to Amyloid β, similar to what was previously reported (*21, 25*). The spatial information, including their distance to Amyloid β, can further confirm their functional and phenotypic diversity. In addition, subcellular mRNA distribution was also informative, showing specific transcript distribution patterns within the cell including polar, random, and centripetal distributions. Extensive changes in these subcellular distributions in glia cells and neurons are found within the Amyloid β microenvironment, most likely caused by changes in the cytoplasm morphology and membrane position, consistent with previous reports (*24*). Cell functions including phagocytosis, stimulus sensing and response, and cell-to-cell interactions are closely linked with these structural alterations during disease development (*23, 24, 26*).

Subcellular mRNA distribution has high heterogeneity across cell types and subtypes. All reactive glia cells were classified as such based on their gene expression, but importantly the mRNA dispersion degree is higher in cells that are closer to plaques within the same type (the larger cytoplasm size is a typical phenotypic feature of glia cell activation in addition to specific gene expression). This new facet of in-situ sequencing data has added one more dimension to the conventional cellular-level gene expression matrix. We assert that a combination of gene expression and subcellular mRNA distribution will improve the understanding of complex transcriptional molecular functions in tissues and lead to more accurate and more quantitative characterizations of cell types and states.

At present, SPRINTseq is yet to be improved in some aspects. Firstly, cell segmentation in brain is a challenge in the field, because cells (including neurons and glia cells) are intricately distributed in the brain, and there is no proper dye that can perfectly define cell boundaries and also is compatible with in-situ sequencing chemistry. Currently we did not take long projection structure (such as axon) into account. One possible improvement is to use expression-guided machine learning approaches for a better cell segmentation. Besides, the scalability holds a great potential to be further improved by engineering optimization. For example, current total time is limited by imaging speed (∼40 min to cover a mouse brain coronal slice) which is incomparable to sequencing reaction (∼15 min for each cycle) in each cycle. Larger FoV lens with lower magnification could reduce the sequencing time spent by half without damaging data quality due to the coding strategy. The current encoding scheme has about 140 original codewords when fewer than 5% of all codeword pairs (including the original codeword and all composite codewords) have Hamming distances less than 3, a preferred condition for error correction. Information acquisition efficiency decreases when more codewords are used to encode a greater number of genes. To encode all 20,000 genes in the human genome, a simple solution is to splice two current 10-bp barcodes into a 20-bp barcode while maintaining the current dilution fold (140^2^ ∼ 20,000). However, when more genes are read, a large dilution fold is needed to maintain crowdedness-robust signal quality, which likely requires a longer barcode.

## Acknowledgments

The authors thank BIOPIC sequencing center for experimental assistant. Funding was provided by National Natural Science Foundation of China Grant 21927802 (Y.H., J.W., P.F.), 22050002 (Y.H.) and T2188102 (Y.H.), Beijing Municipal Science and Technology Commission Grant Z211100003321006 (Y.H.), and Beijing Advanced Innovation Center for Genomics.

## Author contributions

Conceptualization: Y.H. and J.W. Experiment: T.C., W.H., M.J., J.L., J.S., Z.C., Y.P., and G.W. Data analysis: T.C., W.H., M.J., Z.L., M.T., P.F., X.R., and Y.H. Writing: T.C., W.H., M.J., J.W., and Y.H.

## Supplementary Materials in the followed pages

