## Supplementary material for "Rapid and Signal Crowdedness-Robust In-Situ Sequencing through Hybrid Block Coding": Fig. S1

### Materials and Methods

**Cell culture.** The cell lines HEK293T and NIH3T3 were cultured in petri dishes with DMEM (Gibco), 10% FBS (Gibco), and HyClone 1% penicillin-streptomycin solution (Cytiva) at 37°C in a 5% $CO_2$ incubator. To culture cells on a glass slide for imaging, the cells adhering to the dish were treated with 0.25% trypsin and resuspended in culture medium. Meanwhile, a piece of polydimethylsiloxane (PDMS) slab with a hole punched in it was assembled with Polysine slides (Epredia) to create a chamber. The resuspended cells were then seeded into the chamber and incubated for 12–24 hours at 37°C, and then they were washed with PBS buffer (Cellmax) before being fixed.

**Tissue collection and sections.** Animal handling and tissue harvesting methods followed the guidelines and recommendations of local animal protection legislation and were approved by the local committee for ethical experiments on laboratory animals. One 8-week-old wild-type (male), two 10-month-old APP/PS-1 (female) and two 10-month-old wild-type (female) C57BL/6J mice were sacrificed. The brains were harvested, embedded in Tissue-Tek O.C.T. Compound (Sakura), snap frozen, and then kept frozen at -80 °C. The frozen brains were sectioned into 10-μm-thick slices using a CM1950 cryostat (Leica), and each slice was stuck to a Superfrost Plus glass slide (Epredia). The mounted sections were stored at -80 °C.

**Immunofluorescence staining.** Once taken from the -80°C freezer, the brain slices were immediately fixed in 4% paraformaldehyde (PFA) for 15 min at room temperature and then washed in PBS (Cellmax) three times. Next, the slices were dehydrated in a 30% (wt/wt) sucrose solution for 1 hour at room temperature and then washed three times with PBS. The slices were treated with a blocking buffer (5% BSA in PBS-Triton [0.3% Triton X-100 in PBS]) for 1 hour at room temperature and then washed three times with PBS. The primary antibody was diluted in antibody dilution buffer (1% BSA in PBS-Triton) at a recommended ratio of 1:500~1:1000. Next, the slices and the diluted primary antibody were incubated at 4°C overnight and then washed three times with PBS. Finally, the brain slices were incubated with the secondary antibody (1:1000 in antibody dilution buffer) for 1 hour at room temperature (avoiding light), washed three times with PBS, and were ready for imaging.

**Sample pretreatment for sequencing.** Both the cell-on-slide samples and the tissue sections were fixed in 4% (w/v) RNase-free PFA (Shanghai Yuanye) with 0.1% Glutaric dialdehyde at room temperature for 15 min and then washed with PBST (0.25% Tween in 1X PBS). A PDMS chamber assembled onto each slide prevented evaporation while the clonal amplification step was manually performed.

The sample was then permeabilized with 0.01% pepsin in 0.1M HCl at 37°C for 3 min and twice washed with PBST. After that, the samples were dehydrated with 80% ethanol for 10 min and pure ethanol for 2 min and then rehydrated with three PBST washes.

**Probe Design.** A 40nt sliding window was applied to the mRNA transcript sequence of each gene of interest. A bowtie2 (27) alignment of the initial candidate list against the mouse transcriptome was performed to ensure specificity. A successful candidate contained only the original transcript, transcript variants, or predicted transcripts. All probes and other DNA oligomers were ordered from Sangon Biotech (Shanghai). Suitable probe sequence candidates must have had 40%-65% GC content while avoiding five consecutive nucleotides such as 'TTTTT'. The secondary structures of the filtered candidates were further examined and filtered using the OligoMiner (28), and as a final check, a local BLAST query was run on each of the final candidates. The rolling circle amplification (RCA) primer and the sequencing primer were identical sequences on the padlock probes, which each had a unique barcode: three bases were either T or C while the others were G. At each cycle in our 108-probe panel, T and C were evenly distributed. The 'sequencing primer + barcode' block was doubled on each padlock probe to increase the signal-to-background ratio. In our experiment, we tested the various candidates to identify the best probes. For each gene, several (~3) probes with different binding sequences were simultaneously added to the panel. The padlock probes were selected based on their experimental performance in the test. It is also applicable to simultaneously add multiple padlock probes for one gene. All probe sequences in our final 108-gene panel are listed in Extra Table 1.

**Probe hybridization and ligation.** Each sample (fixed and dehydrated) was blocked with oligo-dT (100 nM oligo-dT, 50 mM KCl, 20% formamide, 20 μg/mL BSA, 20 μg/mL Yeast tRNA [AM7119, Invitrogen], and 1U/μL Ribolock RNase inhibitor [Thermo Scientific] in Ampligase buffer [Lucigen]) (29) for 10 min at room temperature. After the samples were placed in a moisture box to prevent liquid evaporation, the probes were hybridized by incubating them in a hybridization mix (200 nM padlock probe for each, 50 mM KCl, 20% formamide, 20 μg/mL BSA, 20 μg/mL Yeast tRNA, and 1U/μL Ribolock RNase inhibitor in Ampligase buffer) at 55°C for 10 min and then 45°C for 1 hour 50 min. After hybridization, the samples were washed with washing buffer (10% formamide, 2X SSC buffer) for three times (10 min per wash) to remove unspecific-binding probes, and then rinsed twice with PBST. Each sample was then incubated in a ligation mix (2.5 U/μL SplintR ligase [New England Biolabs], 20 μg/mL BSA, and 1U/μL Ribolock RNase inhibitor in

SplintR buffer) (29, 30) at 37°C for 2 hours to ligate the nick on the padlock probe, and then the samples were rinsed twice with PBST.

**Amplification and post-fixation.** After padlock probe ligation, RCA was performed using Phi29 polymerase in Phi29 polymerase buffer (New England Biolabs) with 250 μM dNTP (Thermo Scientific), 50 μM aminoallyl-dUTP (Thermo Scientific), 10% glycerol, 20 μg/mL BSA, and 600nM RCA primer at 30 °C for 5 hours, rinsed twice with PBST, incubated with 10 μg/μL $BS(PEG)_9$ (Thermo Scientific) (29, 30) in PBST at room temperature for 15 min to fix the amplified product, and then rinsed three times with PBST. Finally, the sample was washed three times (5 min per wash) with 65% formamide at room temperature and then rinsed twice with PBST.

**Barcode design.** To conveniently locate puncta, a C or a T must have appeared within the first four bases. Also, each barcode should have contained at least one C base to better acquire overall quality sequencing signals. Additionally, to be able to decode co-located barcodes, as well as to reduce optical crowdedness, the Hamming weights were set from 2 to 5, which is an average of 6 G bases across the barcodes. All barcodes that met all those criteria were collected and, based on this library of barcodes, a minimum Hamming distance of 3 between individual barcodes was chosen to improve the orthogonality within the pool. To obtain a library for actual use, an undirected graph of the library was constructed. An edge was added if the Hamming distance between each node in a pair was no less than 3. After the graph was constructed, maximal independent sets (MISs) were calculated from the complement of the original graph. MIS calculations were repeated 10,000 times to reduce the standard deviation of T or C base counts across the final collection (Fig. S9B).

**Flow cell.** Each sample slide was assembled into a flow cell (Fig. 1A) by stacking the slide, a strip of double-sided adhesive tape (20106, ARcare) with a cut-opening that formed a flow channel, and a blank slide with two holes for in-and-out reagent flow. The assembled flow cell was mounted onto our automated imaging/fluidic system (Fig. 1A).

**Fluidics.** An integrated experimental setup that performed the automated sequencing process included fluid delivery, temperature control, motion control, and fluorescence imaging (Fig. S1-2). The system was controlled by LabVIEW software (National Instruments), and a switching valve (C25Z-31814D, VICI) and a Cavro Centris syringe pump (Tecan) were used for fluid selection and fluid volume/speed control, respectively. A pair of two-port solenoid valves

(LVM105RY-5C1U, SMC) were used to construct a bypass for debubbling and saving reagent (Fig.1A, Fig. S1A). The flow cell chip was placed between two home-made manifolds in the fluid system and FEP tubing (I.D. 0.5 mm, O.D. 1/16 inch) connected the pump, manifold, and switching valve. A negative-pressure, fluid drive design prevented reagent cross contamination in the pump.

**Temperature control.** A highly responsive and precise temperature control system was constructed using a Peltier thermoelectric module (Fig. S1A) and a temperature controller (TE Technology). The Peltier module was mounted between the two manifolds in the fluid setup and it closely contacted the bottom surface of the flow cell.

**In-situ sequencing.** To sequence the barcodes corresponding to single molecule RNA, we applied two-color cyclic reversible terminator sequencing-by-synthesis (SBS) chemistry and used NextSeq 500/550 High Output Reagent Cartridge v2 reagents (Illumina) with our modified sequencing protocol (Fig. S2). In brief, the samples were first incubated with sequencing primer hybridization mix (200 nM sequencing primer in 2× SSC and 20% formamide) at 50°C for 3 min and 37°C for 7 min, and then washed with PBS to remove free primers. A complete 15-min sequencing cycle consisting of debubble, cleave, block, incorporate, and deterge. First, 70% ethanol was used to remove possible air bubbles in the flow cell chip, and it was rinsed out with universal sequencing buffer (USB, Illumina). Next, the samples were incubated in cleavage reagent mastermix (CRM, Illumina) at 60 °C for 4 min and washed with cleavage wash mix (Illumina), incubated in 200 mM N-ethylmaleimide (NEM, Illumina) at 60 °C for 2 min, and then washed with USB. Next, the sample was incubated with incorporation mix (Illumina) at 60°C for 160 s and then washed with USB. After incorporation, we washed the sample with 10% SDS and then rinsed it with USB to remove the detergent. Finally, we reduced the temperature to 20 °C and the sample was immersed in universal scan mix (USM, Illumina) for fluorescence imaging. The cleavage step was omitted from the first sequencing cycle but the subsequent cycles used all the steps. In the last cycle, we stained the sample with DAPI solution at 20 °C for 2 min before USM immersion, and that signal was acquired in the last imaging cycle.

**Imaging.** Imaging was performed on a MIM microscopic imaging framework (Applied Scientific Instrumentation) equipped with an S551-2201B motorized stage (Applied Scientific Instrumentation), an ATF6.5 SYS 785 automated focusing module (Wise Device Inc.), a CFI S Plan Fluor ELWD (40X NA 0.60 and 20X, NA 0.70) objective lens (Nikon), an X-Cite Turbo LED light source (Excelitas Technologies), and an Orca Fusion BT scientific CMOS camera (Hamamatsu

Photonics) (Fig. S1A). The following filter sets were used: DAPI-5060C (Semrock) for the DAPI nucleus stain and Cy3/Cy5-2X-B (Semrock) for FISH and the two-color sequencing signal. During each cycle of the sequencing reaction for each mouse brain slice, we imaged ~500 tiles (40X objective lens) with 4% overlap and collected a z-stack of nine planes with a step of ~1.2 μm. The exposure times were 100 ms for Cy3 and 40 ms for Cy5 and DAPI.

**Image processing.** Image processing was performed via Python and MATLAB. Custom codes were used unless references are given. Images from multiple planes were stacked along the z-axis to extend the depth-of-field. Corrected intensity distributions using regularized energy minimization (31) was used to flatten the field of illumination of fused images produced within the same reaction cycle (Fig. S5). To register images acquired in different imaging cycles, the *phase_cross_correlation* function from the scikit-image package (32) was used. A fast Fourier transform was performed to calculate an initial estimate of the pixel shift from the cross-correlation peak for each pair of images and refined to sub-pixel level by a super-sampling factor of 100. Integer shifts were saved to a table and sub-pixel remainders were applied directly to the moving image. In our test, the Cy3 channel was used as the reference channel and all the Cy3 images were aligned with a Cy3 reference image captured from the first sequencing reaction cycle. The same shift was propagated to the Cy5 channel. A global coordinate system was created using the Microscopy Image Stitching Tool (33). The set of Cy3 images within the first cycle was stitched as a reference. The grid positions in the reference cycle, applied with shifts according to registration results, were used to stitch the images from the subsequent reaction cycles. Cell segmentation was performed using adaptive thresholding (*threshold_local* in scikit-image), and cell centroids were located by finding the local maxima after the binarized image was Euclidean transformed. Based on the centroids, the watershed algorithm was applied to further segment the binary mask. The labeled image was finally filtered by setting a minimum cell size.

**Base calling, barcode mapping, and cell segmentation.** After 20 stitched sequencing images had been acquired (10 cycles and 2 channels), the first eight were used for puncta recognition. For each image, we applied white top-hat and then located puncta by finding the image's local maxima (Fig. S6). The extracted coordinates were then used for calling and filtering sequences. The sequences for each punctum were mapped to a library consisting of single and double (composite) barcodes, and before mapping, the library was checked for ambiguous reads. To begin mapping, searches for exact matches for each barcode were done and if no exact match was found, the barcode was compared through the library to find a match with a Hamming distance of 1. Points were dropped as ambiguous barcodes still without matches were found within that tolerance. Mapping information, including position and RNA type, were recorded in a table. Next, we segmented the cells by taking adaptive thresholds using their DAPI-stained images. The binary image then underwent Euclidean transform and was further segmented using the watershed algorithm. After finding each cell's nucleus centroid position, each RNA was assigned to its nearest centroid by constructing a K-D tree. "Cell Index" was added to each RNA in the final readout table and an expression matrix of the cells was generated.

**Spatial co-localization analysis between genes.** A z-score for each gene, across all the cells, was calculated and then analyzed using Pearson's R correlation. A similar analysis was performed on an RNA sub-sample extracted from the read table. For that step, a different sub-sampling factor was used, and an image's binned pixels had a summed value with respect to all its original pixels. Finally, each gene's smaller 2D image was transformed and normalized to a 1D vector, and all the genes' vectors collectively produced a matrix resembling the expression matrix.

**Mapping and annotating cell types.** Several housekeeping and marker genes were used for quality control and to assign major cell types in a divisive manner. For each major type (excitatory neuron, inhibitory neuron, non-neuronal cells), principal components analysis was applied to their normalized cellular expression matrices, and then they were placed in subclusters by using the Louvain shared nearest neighbor (SNN) method. Each subcluster was identified manually by viewing the overall gene expression level. The convex hulls of individual cells were drawn with colors assigned to each cell type, and 2D visualization was done using uniform manifold approximation and projection.

### Supplementary Text

#### 1. Enhancing the signal-to-background ratio

**1.1 Reducing tissue autofluorescence.** One challenge of applying cyclic reversible terminator SBS chemistry (e.g., Illumina sequencing chemistry) to in-situ sequencing is high imaging background noise, which is caused mainly by the non-specific absorption of fluorescent dNTPs by free thiol groups on tissue surfaces. We found that NEM reduces that noise by blocking the free thiol groups, and tissue fluorescent intensity is halved, a significant improvement in the signal-to-background ratio (SBR) (Fig. S3A-B).

**1.2 Number of barcode blocks in a padlock probe.** To either increase the SBR with a constant amplification time or shorten the time needed for RCA while reaching the same SBR, we tested various numbers of barcode blocks, each possessing a barcode and a sequencing primer binding site in a single padlock probe with the same amplification time (enough time). Increasing the number of barcode blocks clearly facilitated the fluorescence SBR (Fig. S3C-F). Two barcode blocks can significantly increase the SBR and ensure a high base calling accuracy. Though three barcode blocks can provide even higher SBR, the signal spot is bigger and will elevate optical crowdedness and affect signal extraction. Therefore, 2x barcode is a balanced choice that can provide sufficient SBR while avoid too much crowdedness.

#### 2. SPRINTseq performance

**2.1 The decoding accuracy of SPRINT-seq.** To verify the decoding accuracy of SPRINT-seq, we used FISH probes to directly hybridize the amplification products, and then we calculated the proportion of FISH counts that co-localized with reads mapped from sequencing to all reads mapped from sequencing (Fig. S7C). The accuracies of probes that targeted different genes or binding sites were greater than 97% (Table S1).

**Table S1. Decoding accuracy validation via fluorescence in-situ hybridization (FISH)**

| Probe | Read Counts | FISH Verified | Decoding Accuracy |
|---|---|---|---|
| **Actb_pd1** | 12658 | 12218 | 97% |
| **Actb_pd2** | 16715 | 16405 | 98% |
| **Actb_pd3** | 10718 | 10428 | 97% |
| **Slc17a7_pd1** | 25521 | 24883 | 98% |

| Probe | Read Counts | FISH Verified | Decoding Accuracy |
|---|---|---|---|
| **Slc17a7_pd2** | 22944 | 22506 | 98% |
| **Slc17a7_pd3** | 14500 | 14105 | 97% |

**2.2 The sensitivity of SPRINT-seq.** The number of Actb sequencing signals (one probe per gene) that we counted in sampled cultured cells averaged 132 spots per cell (Fig. S7D-E). Then we compared that result with the result from a QX200 digital PCR system (Bio-RAD), which was 347 signals per cell on average. This shows that the probes captured more than 30% of the target mRNAs (Fig. S7F). The overall accuracy and sensitivity of SPRINTseq was tested through comparison with RNAscope profiling of the same gene (Actb) in adjacent slice (within 50 μm) in the same brain region from the same mouse. The gene pattern is similar and average counts per cell is comparable (93.6 cp v.s. 96.7 cp, Fig. S7G).

**2.3 Cyclic attenuation of SPRINT-seq signals.** As with any non-single molecule sequencing chemistry, reaction signals will gradually decay along with the reaction cycles in SPRINT-seq. The signal is proportional to the nucleotides that were incorporated into the nascent strand. Commercially available Illumina sequencing reagent kits perform with a small-decay rate so they can generate hundreds of bases per sequencing run, more than enough to sequence 10–20 nt barcodes for the SPRINTseq approach. Our experimental results showed a small signal decay through the cyclic reaction (Fig. S8), and such a small signal attenuation should not impair base calling accuracy.

### 3. SPRINT-seq coding design

**3.1 Hybrid block code. I**n coding theory, a block code corresponds to an injective mapping,

$$C: \Sigma^k \to \Sigma^n$$

Here, $\Sigma$ is a finite and nonempty set, which is denoted as the alphabet. The message $m$ and the codeword $c = C(m)$ are elements of $\Sigma^k$ and $\Sigma^n$, respectively, and $k$ and $n$ are the message length and block length, respectively. Both $k$ and $n$ are positive integers. Typical binary block code has an alphabet satisfying $|\Sigma| = 2$, which indicates 2 possible letters in a message or codeword.

There are finite types of amplicons in a single in-situ sequencing experiment. Thus, a bijection can be formed between amplicon type and a binary string. Without loss of generality, a set of 7-letter strings can represent 128 distinct types of amplicons, which is greater than the 108 types of transcripts we later chose. Based on this bijection, each message can be transformed into a codeword with length $n = 20$.

To reduce decoding time, each of the 10 reaction rounds received two codewords in parallel, and they can be combined into an $n = 20$ codeword. The combination of the two $n = 10$ codewords is not a direct product, but rather has several constraints. For example, the A nucleotide from an Illumina 2-channel SBS had two types, each with different fluorescent dyes (Fig. S9A), so signal recognition and error correction were harder. Thus, the A nucleotides were not used for barcoding. This rule added a constraint between the two codewords, preventing "1" bits from occurring at the same sequential position. For instance, the codeword combination "0100010010" and "0100001001" is not permitted because of the coexisting signal at the second bit. This specific coding scheme is named 'hybrid block code'.

While a total of $3^{10} = 59{,}049$ barcodes were enumerated in the initial collection, a Hamming weight range between 2 and 5 was chosen to avoid an excessive use of each bit position. In addition, at least a C or a T base within the first four bits was used as a filter condition, thus making the locating of initial puncta more robust. One or more C bases are required in the barcode for a better overall signal quality. These rules yield a library of 11,341 barcodes. Also, the minimum distance $d$ of the block code should be large enough to allow error detection and correction. The minimum distance of the code $C$ is defined as

$$d := \min_{\substack{m_1, m_2 \in \Sigma^k \\ m_1 \neq m_2}} \Delta[C(m_1), C(m_2)].$$

For two codewords, $\Delta(c_1, c_2)$ denotes the Hamming distance between $c_1$ and $c_2$. As a demonstration, $d = 3$ was chosen. To obtain a subset of the library that satisfies this constraint, an undirected graph $G = (V, E)$ was constructed using all the barcode sequences in the library. $V$ was the set consisting of all the nodes, and $K$ denoted all the 2-element subsets of $V$. $E$ is a subset of $K$. Two nodes in the graph are connected by an edge if the Hamming distance between them is no less than 3 ($\Delta(c_1, c_2) \geq 3$). After 10,000 trials, the MIS of the complement graph was $H = (V, K \backslash E)$. The MIS with the most uniform signal distribution was selected as a final barcode list (Fig. S9B).

To evaluate the sequencing signal distribution, a variance was calculated from the signal counts of each bit within the library, so given a list $B = \{c_1, c_2, \dots, c_N\}$ with $N$ barcodes, where $c_1(i)$ denotes the $i$-th letter of codeword $c_1$ (0 or 1), the variance $\sigma^2$ is calculated as

$$\sigma^2 = \frac{1}{20}\sum_{i=1}^{20}\left(\sum_{j=1}^{N} c_j\,(i) - \mu\right)^2,\ \mu = \frac{1}{20}\sum_{i=1}^{20}\sum_{j=1}^{N} c_j\,(i).$$

For the hybrid block code mentioned above, such a metric can be calculated by counting T and C basewise across the library. A final collection of 369 barcodes (Extra Table 2) was produced under this criterion, with $\sigma^2 = 2.3475$ and a mean Hamming weight of 4.7669. In each round, T and C were evenly distributed (Fig. S9D).

**3.2 Barcode orthogonality.** As previously described, the hybrid block code has a minimum distance of $d = 3$. This coding enables error detection and correction between individual amplicons. However, as the signal density increases, two proximal RCA products have a significant probability of being co-located in the image and causing signal overlap (polyclone). This phenomenon poses a challenge to the decoding process.

To circumvent this problem, we generated a set of superimposed barcodes (composite codewords) that were combinations of a pair of barcodes in the library. For example, the combination of “GCTCGGCTGG” and “GCTGGTGCGC” yields “GCTCGTCAGC” and represents signal overlapping. These new sets of barcodes were merged with the original library. The Hamming distances between each single or overlapping barcode pair were calculated and if any two of the barcodes were the same ($\Delta(c_1, c_2) = 0$), they would both be added to the ambiguous barcode set. This set was then removed from the merged library and the rest of the barcodes were checked again for ambiguity.

Among the 6,105 barcodes $[110 + \binom{110}{2}]$, 234 ambiguity-prone barcodes were eliminated. This modified reference for amplicon type calling ignores a slight amount of sequencing signals but ensures robust identification of most of the barcodes. To characterize the orthogonality, the Hamming distances of the 17,231,385 codeword pairs, including single and superimposed barcodes, were collected and only a few pairs (0.199% of all pairs) could cause ambiguous decoding (Fig. S9E).

**3.3 Encoding capacity.** Encoding capacity is a metric that assesses the general scalability of the coding design. Each in-situ detection method, both hybridization and sequencing based, has its characteristic encoding capacity that can be summarized in a few key parameters (Table S2).

SPRINT-seq achieves a high encoding word count using only 20 bits. However, compared to its FISH-based counterparts, this encoding capacity carries the cost of a higher Hamming weight (an average of 7). Notably, the process that MERFISH, seqFISH+, and SPRINTseq used to obtain ‘bits for decoding’ differed from that used by STARmap and BARseq2, which instead of thresholding, use vector projection for decoding or quality filtering without redundant bases. Thus, the bits for the decoding parameter cannot be directly compared.

**Table S2. Encoding capacity comparisons of spatial in-situ detection methods.**

| Method | Reaction Cycles | Imaging Channels | Bits for Decoding | Hamming Weight (mean) | Distinct Codewords |
|---|---|---|---|---|---|
| **MERFISH** | 23 | 3 | 69 | 4 | 10050 |
| **seqFISH+** | 80 | 1 (3) | 80 (240) | 4 | 3333 (10000) |
| **STARmap** | 6 | 4 | 24 | Not defined | 1020 |
| **BARseq2** | 7 | 4 | 28 | Not defined | 65 |
| **SPRINTseq** | 10 | 2 | 20 | 4-10 (7.073) | 5871 |

Since only part of the initial barcode library was used for the experiment, we had to validate whether the overall encoding performance of the whole library resembled that of its subset. As mentioned previously, some overlapping barcode pairs were removed from the reference library to avoid ambiguity. In general, fewer percentages of those overlapping pairs can be used for error-correcting decoding references as the barcode library is used more (Fig. S9F).

To control the loss from identified overlapping barcodes, the number of used barcodes must be restricted. For example, for an acceptable drop rate (0.05), 140 barcodes out of 369 is the maximal subset size. This subset generates 9,393 codewords that can be encoded with 13.2 binary bits. The 140-barcode subset yields a rate ($R = k/n$) of 0.660, higher than that from a Hamming(7,4) code (0.571) but lower than that of a Hamming(15,11) code (0.733). This demonstrates the difficulty of designing a practical error correction code with a higher rate, given $n = 10$. Considering the basic code scalability, it would require no less than 40 bits of information to encode amplicons at the transcriptome level ($140^2$, about 20,000 types of RNA). Two orthogonal rounds of barcode calling should achieve the encoding scheme, but the optical crowdedness issue prevails over the problem of encoding capacity, and will be discussed later.

**3.4 Information entropy and encoding efficiency.** A rough estimation of the encoding efficiency can be demonstrated. Assuming the amount of each amplicon type is uniformly distributed, a punctum $X$ in an arbitrary field of view is equally probable ($p$) to be any of the amplicon types. That is, for any barcode $c_i$ in the library $B = \{c_1, c_2, \dots, c_N\}$, we have

$$P(X = c_i) = \frac{1}{N} = p.$$

Without observation, the barcode has an initial information entropy ($H$) of

$$H(X) = -\sum_{i=1}^{N} P\,(X = c_i)\log P(X = c_i) = -\log p = \log N.$$

Encoding efficiency can be characterized by calculating the average entropy decrease per reaction cycle $\Delta H_{\text{cyc}}$ and average entropy decrease per imaging round $\Delta H_{\text{im}}$ (Table S3).

**Table S3. Encoding efficiency comparison**

| | Reaction Cycles | Imaging Channels | Distinct Codewords | $H(X)$ | $-\Delta H_{\text{cyc}}$ | $-\Delta H_{\text{im}}$ |
|---|---|---|---|---|---|---|
| **MERFISH** | 23 | 3 | 10050 | 4.002 | 0.174 | 0.058 |
| **seqFISH+** | 80 | 1 | 3333 | 3.523 | 0.044 | 0.044 |
| **STARmap** | 6 | 4 | 1020 | 3.009 | 0.501 | 0.125 |
| **BARseq2** | 7 | 4 | 65 | 1.813 | 0.259 | 0.065 |
| **SPRINTseq** | 10 | 2 | 5871 | 3.769 | 0.377 | 0.188 |

Notes: $H(X)$, initial information entropy of a given barcode; $-\Delta H_{cyc}$, average entropy decrease per reaction cycle; $-\Delta H_{im}$, average entropy decrease per imaging round.

When SPRINT-seq is significantly more efficient than other methods, $-\Delta H_{\text{im}}$ indicates the average information per observation (imaging) for a sequenced amplicon. This property also greatly contributes to imaging time efficiency.

**3.5 Error detection and correction.** The orthogonality of SPRINTseq encoding makes error correction possible in in-situ sequencing (Table S4).

**Table S4. Error detection and correction comparison**

| | Encoding Bits (Binary) | $d$ (Binary or Quaternary) | Error Detection | Error Correction |
|---|---|---|---|---|
| **MERFISH** | 69 | 4 | Yes | Yes |
| **seqFISH+** | 80 | 2 | Yes | Yes |
| **STARmap** | 24 | 1 | Yes | No |
| **BARseq2** | 28 | 3 | Yes | No |
| **SPRINTseq** | 20 | 3 | Yes | Yes |

Note: *d*, minimum distance of a block code.

Furthermore, this correction can deal with a polyclonal signal, which is a common issue during high-throughput detection.

### 4. Signal dilution in relieving crowdedness

**4.1 Theoretical count limit under crowdedness.** The physical crowdedness is mainly caused by DNA nanoballs, amplified through RCA. The diameter of an amplified DNA nanoball is roughly 300~400 nm. If we simply calculate the theoretical count limit based on volume, the physical environment of a cell (d ~ 10 μm) can accommodate more than 20,000 DNA nanoballs. While with the current imaging system, only a few hundred to one thousand spots could be resolved per round per channel. And this limit under optical condition could be potentially improved by increasing physical dilution (selective amplification) and optical dilution ratio and increasing signal extraction strategy.

**4.2 Selective amplification.** To validate the effect of selective amplification, we performed FISH experiments on two RCA amplified brain slices. To begin, 20% functioning padlock probes were added to a “masked” group while the control group had no irreplicable probes. As a measure for effective imaging area, Actb was fully amplified in both mask and control groups. An 800x800 $μm^2$ region from each slice was arbitrarily chosen. After locating the signals’ positions, neighbor pairs within a distance threshold $l$ were counted and were denoted as $M$. To characterize signal density independent of the total signal count $N$, the average neighbor pair count was introduced as $m = M/N$.

Since we are interested in the density difference, we considered a simple situation in which signals are uniformly distributed across the field of interest. Let $S$ be the total area, then $\sigma = N/S$ denotes the constant signal density. Given a central position and a search radius $r$, the average neighbor count $m$ behaves quadratic with respect to $r$:

$$m = a\sigma r^2,$$

where $a$ is an arbitrary constant. For two samples with different signal densities, $\sigma_1$ and $\sigma_2$, we have

$$\frac{m_1}{m_2} = \frac{\sigma_1}{\sigma_2}.$$

This indicates that the neighbor count can characterize signal density. However, the actual distribution is much different than that of this toy model, and the two distributions approach equality only when $l$ is significantly large. Nevertheless, the quantity $m_1/m_2$ can still be calculated, and we named it the amplification dilution factor. To obtain an accurate dilution factor, cross-sample correction is needed because mask and control experiments may exhibit different amplification efficiencies or other experimental differences (Fig. S10A-B).

Before cross-sample correction, the larger amount of Actb amplicons in the control group caused a decrease in calculated values compared to those after correction (Fig. S10C-D). The dilution factor of other genes, such as Snap25, was thus divided by $N_1/N_2$ to compensate that distortion. The corrected dilution factor was close to our expected factor of 0.2, indicating that the quantity can be accurately re-calibrated (Fig. S10D). The observed amplification dilution factor is higher than expected when the distance is small, mainly because the average neighbor signal number is small (less than 5) in signal-sparse areas at small distances. Thus, the calculated dilution fold is higher than 0.2. The dilution fold will converge to a true density ratio as the distance becomes longer. This process fits well with exponential decay (Fig. S10E).

To determine which genes to be masked, highly expressed genes are identified through single-cell expression data sets, which were generated by many scientists recently, and can be easily found in the public domain. The optimal dilution ratio can be adjusted through multiple experiments, while in most cases this factor can be determined empirically. Genes that are highly expressed in all cells (e.g., Actb) or expressed in a few cells but with a high density (e.g., Sst) were chosen as the genes to be masked. Typically, we will let the spot count from one gene within one cell to be less than 30 to avoid severe physical crowdedness. We found for most cases a dilution ratio around 5 is sufficient to avoid most crowdedness issues.

**4.3 Optical dilution.** To achieve greater possible throughput, the fluorescent signals should be dispersed by adding imaging channels or reaction cycles, and this is generally described as 'optical dilution' (Fig. S10F). Given an imaging channel count $N_{\text{chn}}$, total reaction cycle count $R_{\text{cyc}}$, and signal count $w$ (Hamming weight of a single type of RNA/amplicon), we have the optical dilution fold as

$F_{\text{dilute}} = \frac{R_{\text{cyc}}}{w} N_{\text{chn}}$ (optical dilution factor).

SPRINT-seq offers greater flexibility with greater optical dilution fold than most other in-situ sequencing-based approaches which are not easily scaled up because they lack a silent channel (Table S5).

**Table S5. Optical dilution comparison**

| | Imaging Channels | Reaction Cycles | Signal Counts | Optical Dilution Fold |
|---|---|---|---|---|
| **MERFISH** | 3 | 23 | 4 | 17.25 |
| **seqFISH+** | 1 (3) | 80 | 4 | 20 |
| **STARmap** | 4 | 6 | 6 | 4 |
| **BARseq2** | 4 | 7 | 7 | 4 |
| **SPRINTseq** | 2 | 10 | 2-5 (4.77) | 4.20 |
| **SPRINTseq (potential)** | 3 | 20-30 | 6-9 | 10 |

As mentioned in Section 3.3, to scale up in-situ sequencing toward the transcriptome level, our method would require two sets of current 20-bit block code, which is 20 reaction cycles and an average $w$ of 9.534. However, with such a Hamming weight, the reaction cycle count would have to be 96 with two channels and 64 with three channels to satisfy an optical dilution fold of 20. Therefore, other methods, such as selective amplification and expansion microscopy, can be combined to meet the dilution requirement and thus increase time efficiency. We estimate 20–30 cycles of reaction may sufficiently cover 20,000 types of transcripts with three-channel imaging (optical dilution fold of 10 and overall dilution fold of 20). Even with two-channel imaging, which is compatible with commercial 2-color sequencing reagents, 50 reaction cycles should be a sufficient cover.

### 5. SPRINTseq of mouse brain slices

**5.1 Raw sequencing images.** Four slices of whole brain (from 4 mouses, 2 normal mouse and 2 mouse with Alzheimer's disease) were prepared for sequencing at the same time, and each prepared slice was sequencing sequentially (4 x 9.5 hours because we used a 40x microscope objective lens, with a 20x microscope objective lens it will take 4 x 5 hours in this step). The whole process from sample to data takes 48 hours (Fig. S11A).

The whole brain (about $10.2 \times 7.6\ \mathrm{mm}^2$) was divided into about 500 (513 for normal brain shown in Fig. 3 and 462 for the Alzheimer's disease [AD] brain shown in Fig. 4) field-of-view (FoV) tiles and at nine focal planes per FoV. Each image contained $2304 \times 2304\ \mathrm{px}^2$, equivalent to $374.4 \times 374.4\ \mu\mathrm{m}^2$. For the experiment, we took 10 cycles, 2 channels per cycle, of sequencing images with a final DAPI channel image to identify nuclei (Fig. S11B). We ultimately captured approximately 100,000 images, which is equal to 530 billion pixels. There were four gene categories containing 108 genes in the panel for profiling: (1) house-keeping genes, (2) marker genes that targeted each cell type, (3) genes that have distinct spatial-expression patterns in specific regions, and (4) genes that are potentially responsive in AD. Additionally, two genes (A and B) were added as negative-controls to check the false-positive rate. Negative-control gene A (human BMP4) is not expressed in mouse brain, so it was used to check the false probe-binding (and ligation) rate. We obtained 7,328 reads for gene A, which was equal to 0.05 reads per cell. Through transversion of its primer binding sequence, the padlock probe of negative-control gene B (human Sox2) was designed to be irreplicable so it could test the false barcode mapping rate. We obtained 3,224 reads for gene B, which was equal to 0.02 reads per cell. As a reference, we got 1,668,416 reads for Actb after 0.2x selective amplification, and that was equal to an average of 57.08 reads per cell.

**5.2 Whole brain cell classification.** We used the normal mouse brain section shown in Fig. 3 to illustrate our cell classification strategy (Fig. 3, Fig. S12). First, we performed quality control on all cells in the expression matrix. Since the thickness of a brain slice was 10 μm, some cells may not have been morphologically "complete", so only a small fraction of them were present on the slice and that resulted in a low read number for such cells. The expression levels of certain housekeeping genes were chosen as the threshold. Actb expression exceeded 20 copies in 123,491 (84.5%) of all 146,137 cells, and those cells that passed primary filtering were used in the following analysis. It is difficult to accurately classify the cell types of all 123,491 cells, so we divided all cells into three general categories based on marker gene expressions: excitatory neurons (Slc17a7+; 40,753 cells), inhibitory neurons (Gad1+/Gad2+; 6,178 cells), and non-neuronal cells (76,560 cells). To make classification easier, cells of similar types should be classified into the same category prior to subsequent analysis.

For each category, we first performed a preliminary classification through Louvain SNN. A few cells expressed in an unexpected manner. For example, some excitatory neurons (Slc17a7+) also expressed inhibitory neuron marker or neurotransmitter genes (e.g., Vip). Although we could not rule out the possibility that these cells might truly exist in an evolutionarily ancient brain region, very likely it may have been caused by imperfect RNA assignment in cell-dense regions. Thus, we dropped the cells that had a highly unreasonable expression pattern and then we performed a second cell classification for each category. Ultimately, we used 39 genes to identify 17 cell types with distinct expression patterns among the remaining 113,625 cells (92.0% of 123,491 cells; Fig. S12B). We compared the expression pattern in each annotated cluster against single cell sequencing (combined by cortex, hippocampus, thalamus and hypothalamus data, from http://mousebrain.org) result. The clusters in SPRINTseq have a generally good correlation with their corresponding clusters in single cell sequencing data. (Fig. S13)

**5.3 Comparison of AD and normal mouse brain in gene expression.** We first compared the total read numbers between each gene of the AD and normal mouse. The expression levels of the target genes at the whole slice scale did not change significantly in AD mice (Fig. S15A). Similarly, when we examined different brain regions, those genes' expression levels between normal and AD mice were also comparable. However, some genes' spatial distributions changed significantly, and the subsequent spatial correlation analyses showed that they were enriched with amyloid plaques (Fig. 4A-B, Fig. S15B). This demonstrates how important spatial information is as another dimension independent of expression level information.

### 6. Improving SPRINTseq throughput by using low magnification lens (20x)

Current imaging step takes ~2/3 of total sequencing time (~40min) for mouse brain coronal slice in SPRINTseq. The throughput can be further improved by shortening imaging time if 20x lens instead of 40x are used. 40x or higher magnification lens are usually used for signal spots imaging in in-situ sequencing because of low signal intensity, signal crowdedness and the quest for higher-resolution. Due to the high signal intensity and the coding strategy that allows us to accurately locate the exact signal position from crowding space, SPRINTseq can overcome the issues from low magnification and use 20x lens to further improve the throughput.

We used 20x and 40x lens to image the same sequencing cycle (Cycle 1, Cy5-channel) to directly compare their imaging quality (Fig. S17A). Although the 20X images are more blurred, the high signal-to-background ratio still make the spot identification feasible and our data show that the identification result is almost identical to that from 40X images. Then we compared gene distribution pattern obtained from sequencing results imaged by 20X lens with those by 40X lens (Fig. S17B-C. The data are obtained in different experiment, but brain slices are adjacent in 100 μm). The spearman correlation r of 110-gene expression level (log2(counts)) between them reached 0.968, which is similar to correlation between 40x and 40x (r = 0.972).

In a nutshell, both 20x and 40x lens are acceptable for providing highly accurate and precise results, and the data quality is similar. 20x lens takes 5 hours for sequencing mouse brain coronal slice while 40x lens takes 9.5 hours.

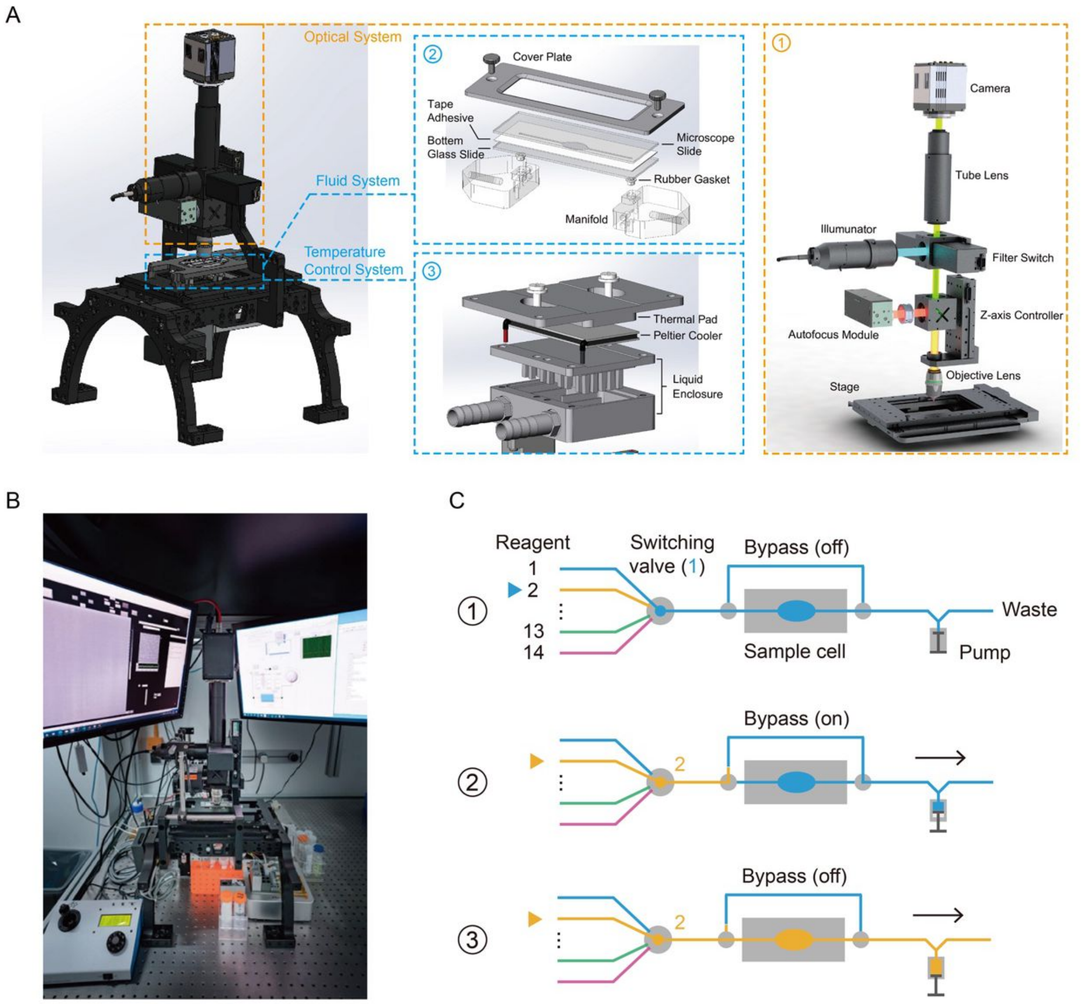


**Fig. S1. Setup construction and automation. (A)** The setup can be divided into three major subsystems: optical, fluid, and thermal. Silicone grease was added to both sides of the Peltier cooler for better thermal conductivity. A rubber gasket between the upper and lower liquid enclosures sealed them together. **(B)** Photo of a functional setup fixed on an optical table. **(C)** Diagram of the fluidic exchange process. A bypass pipeline was used to preload the reagents to avoid inconvenient loading caused by dead volume. A small volume of air was inserted between two distinct reagents to avoid mixing. To load sequencing reagents, we turned the switching valve to the next-round reagent pipeline when the last part of its volume had been loaded into the flow cell. In this way, the reagent that was to be loaded in the next round had already filled up the pipeline's "dead volume" in front of the flow cell, thus efficiently saving both time and cost.

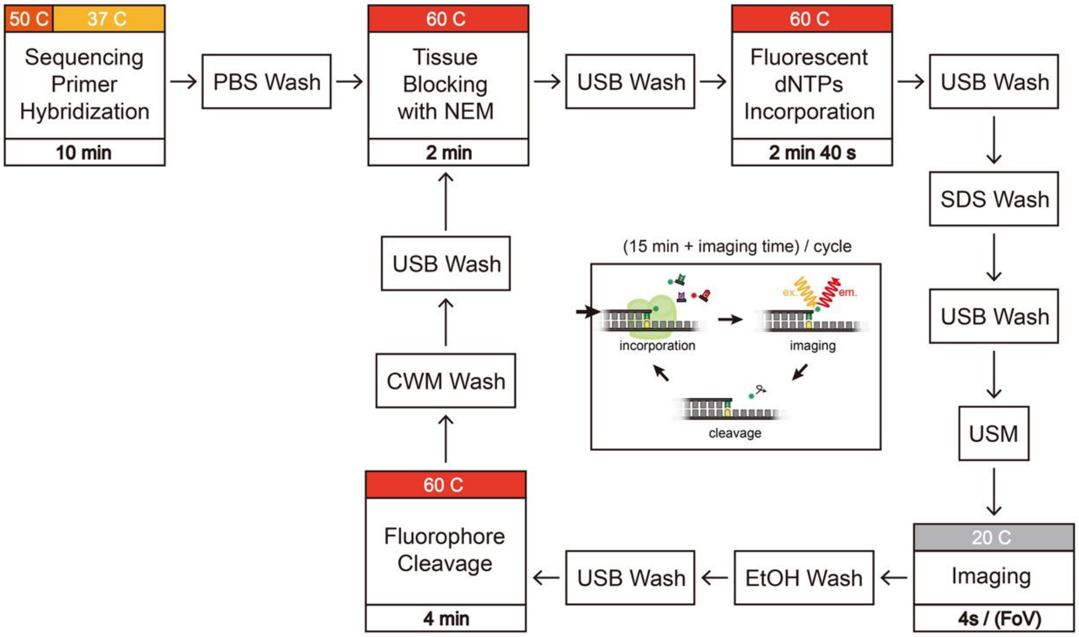


**Fig. S2. Block diagram of the sequencing process using Illumina sequencing-by-synthesis reagents.** Each cycle takes about 15 minutes per reaction. The main step includes tissue blocking, fluorescent dNTP incorporation, and fluorophore cleavage. For a 9-plane z-scan imaging program, each field of view (FoV) took 4 seconds. NEM: N-ethylmaleimide, USB: universal sequencing buffer, USM: universal scan mix, CWM: cleavage wash mix.

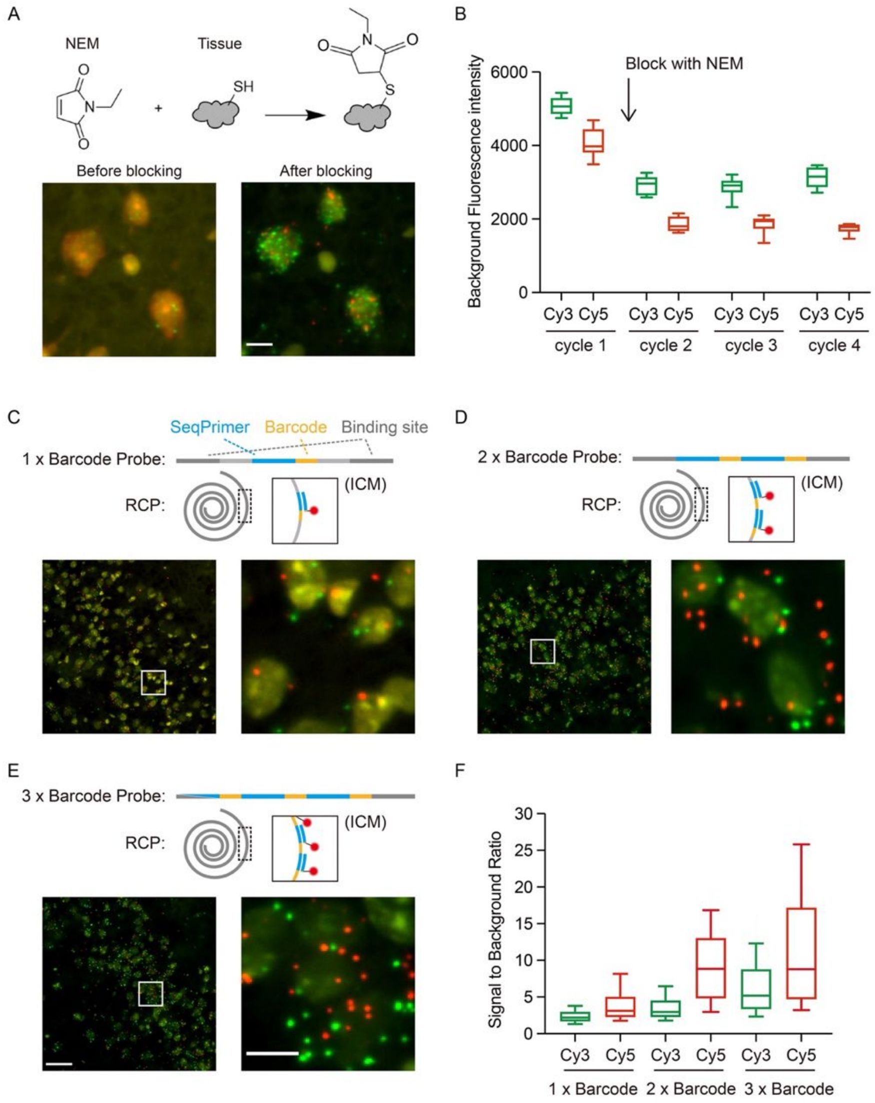


**Fig. S3. Improvement of signal-to-background ratio. (A-B)** Tissue auto-fluorescence reduction via N-ethylmaleimide (NEM) blocking. By preventing fluorescent dNTP absorption by free thiol groups (SH), NEM blocking reduced the average background intensity by about 40%. Photos show the first cycle (Before blocking) and the second cycle (After blocking). Scale bar: 10 μm. **(C-F)** Signal-to-background ratio (SBR) improvement through a multi-barcode design. **(C)** The barcode block on a padlock probe was **(D)** doubled to increase, or to attain a similar, SBR within a shorter amplification time. **(E)** A 3x barcode was also tested and quantitatively compared with mono and double ones. RCP: rolling circle product, ICM: incorporation mix. **(F)** The 2x barcode was sufficient for signal calling so it was adopted. Scale bar: 50 μm for bigger field and 10 μm for the enlarged image. In the graphs, the boxes show the interquartile range, the lines in the boxes are the medians, and the bars show min/max values. Cy3 and Cy5 are base-dependent fluorescent dyes for T and C respectively.

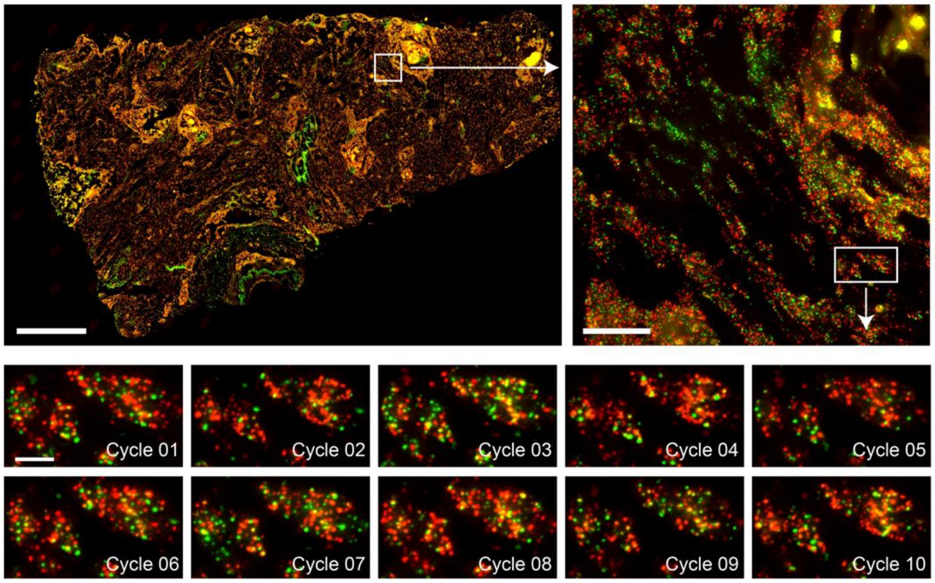


**Fig. S4. Sequencing performance on high-fluorescence-background tissue.** SPRINTseq raw image results on human oral squamous cell carcinoma (OSCC) sample. Scale bar: 1 mm upper left, 50 μm upper right, 10 μm below.

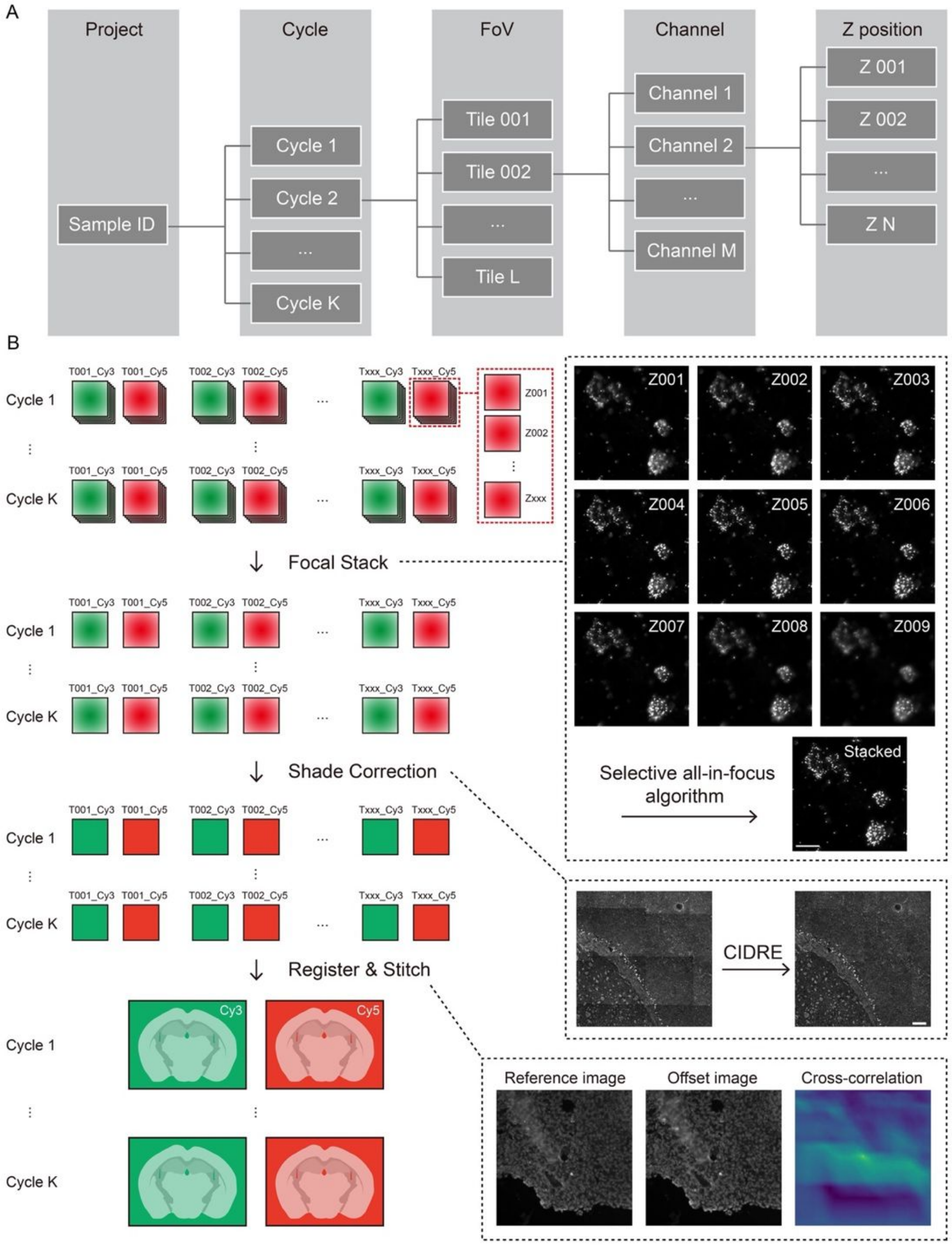


**Fig. S5. Image correction and alignment. (A)** The structure of our data. FoV, imaging field of view. **(B)** Processing from raw images to final images. Images with different z-axis positions were first stacked into one image for each tile (T, Scale bar: 10 μm). Then, images were captured using CIDRE shade correction (Corrected Intensity Distributions using Regularized Energy minimization) to flatten the field of illumination (Scale bar: 100 μm). All images from 10 cycles in 2 channels were registered tile by tile through maximizing 2-D cross-correlation, using the images of the Cy3 channel of Cycle1 as a reference image. Finally, images from 10 cycles in 2 channels (Cy3 and Cy5) were stitched together to generate 20 whole brain signal distribution images.

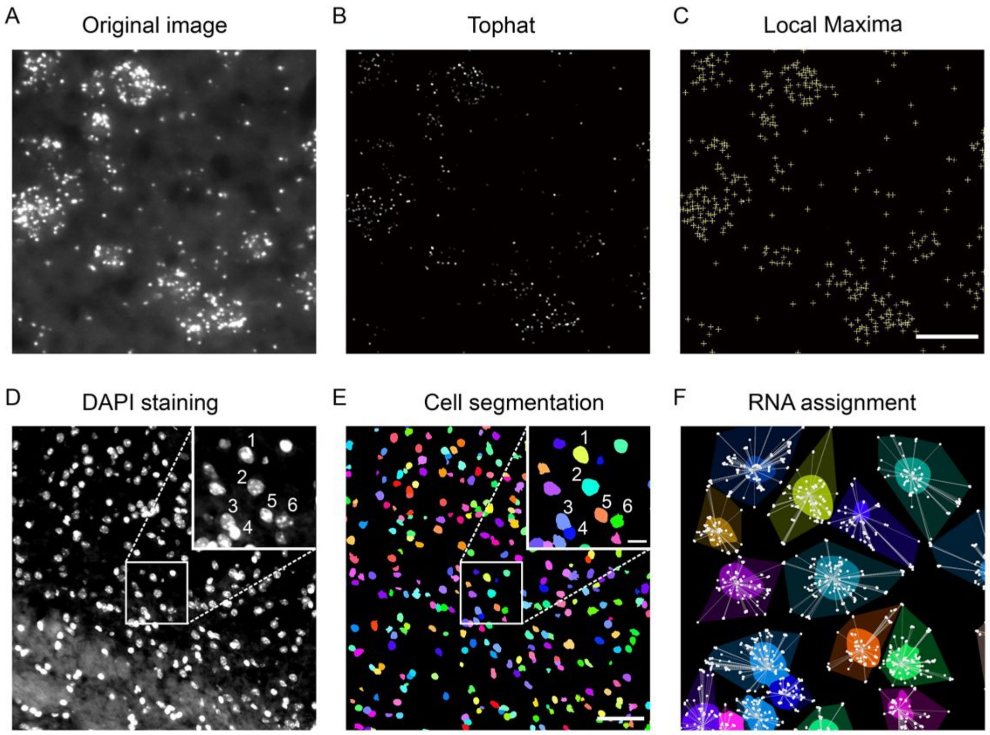


**Fig. S6. Signal calling, cell segmentation, and RNA assignment. (A-C)** Signal calling. After applying the white tophat transform, each punctum was located by finding the local maxima over the image. Scale bar: 10 µm. **(D-E)** Cell segmentation. With the cell nuclei stained with DAPI, the cells were segmented by taking adaptive thresholds. The binary image then underwent Euclidean transform and was further segmented using the watershed algorithm. Scale bar: 100 µm, large field; 10 µm, enlarged inset. **(F)** RNA assignment to segmented cells in **(E)**. Nucleus centroid positions were found and a K-D tree was constructed to assign RNA reads to the nearest centroid. When RNA was determined by signals from multiple cycles (i.e., in sequencing cases), each RNA was first identified through base calling and barcode mapping before they were assigned

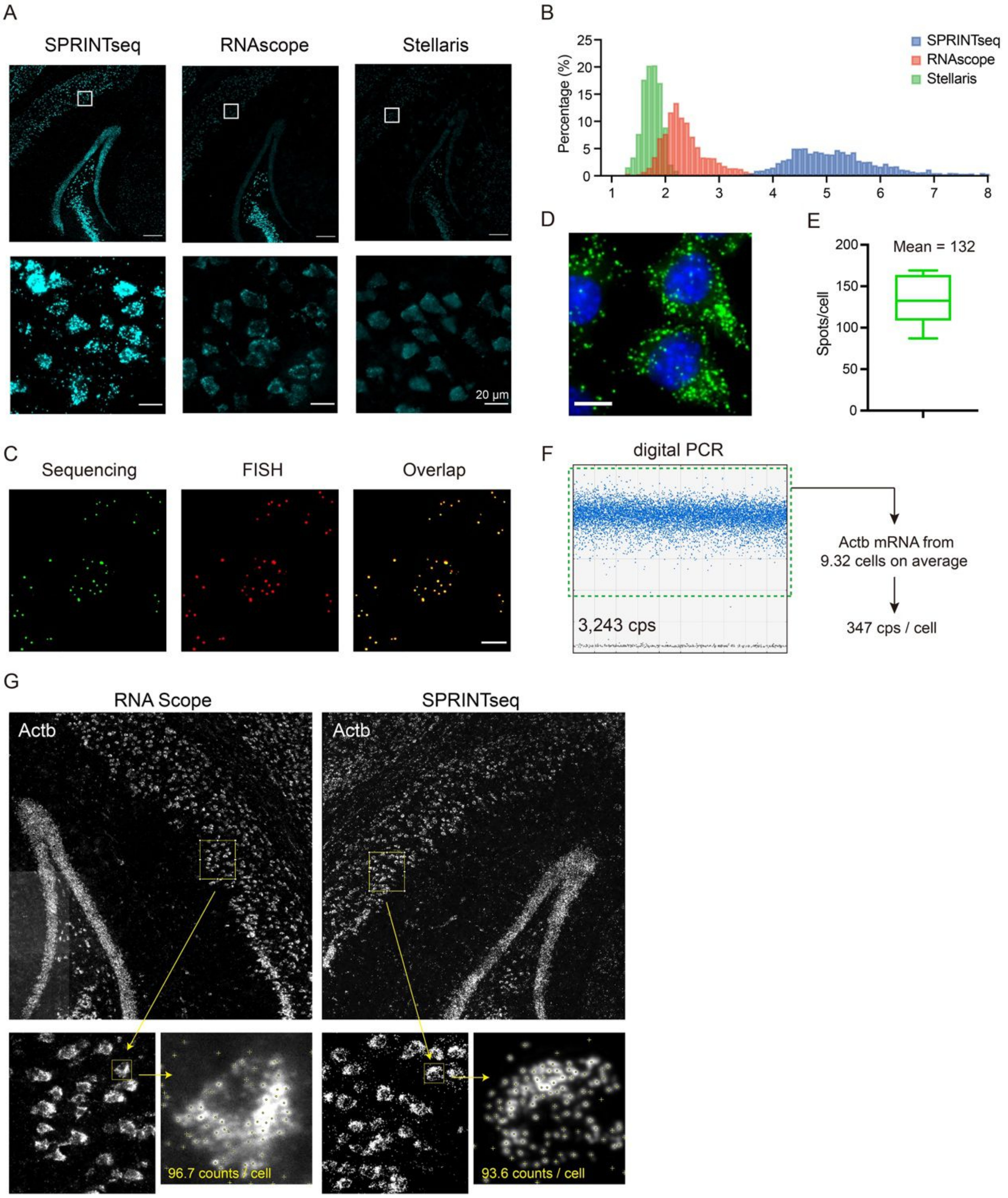


**Fig. S7. Sequencing characterization: SBR, accuracy, and sensitivity. (A)** Photos showing the high SBR of SPRINTseq compared to those of commercial technologies (RNAscope [ACD Bio] in-situ hybridization [ISH] and single-molecule Stellaris [Biosearch Technologies] fluorescence ISH [smFISH]). Results are from within the same region of mouse brain and were captured with an epi-fluorescent microscope, 40 x objective lens, and a sCMOS camera. The white box areas in the top photos are enlarged in the bottom photos. Scale bar: 200 µm above, 20 µm below. **(B)** Quantitative comparison of SBR distributions in **(A)** show that SPRINTseq had the highest SBR. **(C)** Decoding accuracy verification through

FISH after sequencing. FISH probes were used to directly hybridize the amplification products after sequencing (10-nt barcode), and the proportion of co-localized reads were calculated. More than 97% of sequencing reads overlapped with FISH reads, indicating a high accuracy of base calling and barcode mapping. Scale bar: 10 μm. **(D-E)** Sensitivity of SPRINTseq characterization in mouse 3T3 cells. An average of 347 reads (gene: Actb) per cell were obtained through digital PCR (Bio-RAD model QX200) and 132 reads per cell, on average, were obtained through SPRINTseq. The resulting sensitivity was about 38%. Scale bar: 10 μm. The box in the graph shows the interquartile range, the line in a box is the median, and the bars show min/max values. **(F)** Digital PCR quantification of Actb mRNA in mouse 3T3 cells. The mRNA was extracted from ~116,590 cells and reverse transcribed. After 1000x dilution, a 4 μl sample was spiked into 46 μl digital PCR master mixes. The green dotted box encloses positive signals. The mean total positive signal of three replicate tests was 3,243 copies (cps). **(G)** Overall accuracy through cross-methods comparison. The gene distribution of Actb by SPRINTseq (1 probe) and RNAscope respectively in adjacent slice (within 50μm distance) in the same brain region from the same mouse. The gene pattern is similar and overall sensitivity is comparable.

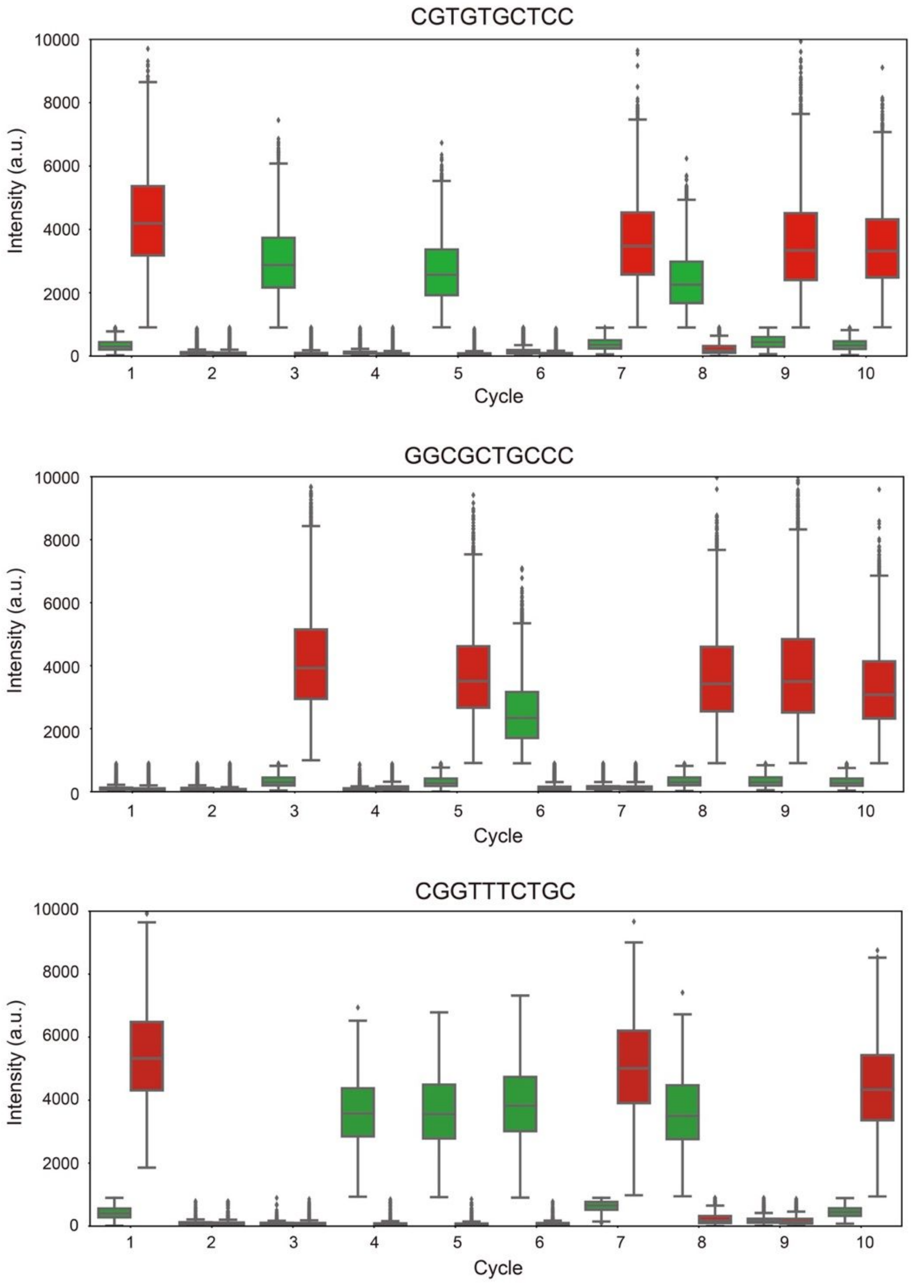


**Fig. S8. Signal intensity through cyclic sequencing.** Signal intensities of three sample sequences. No obvious dephasing was observed, and cyclic signal attenuation (for both colors), in terms of base calling, was negligible. Red represents Cy5 labeled 'C' and green represents Cy3 labeled 'T'. The boxes show the interquartile range, the line in a box is the median, the bars show min/max values, and the dots are outliers.

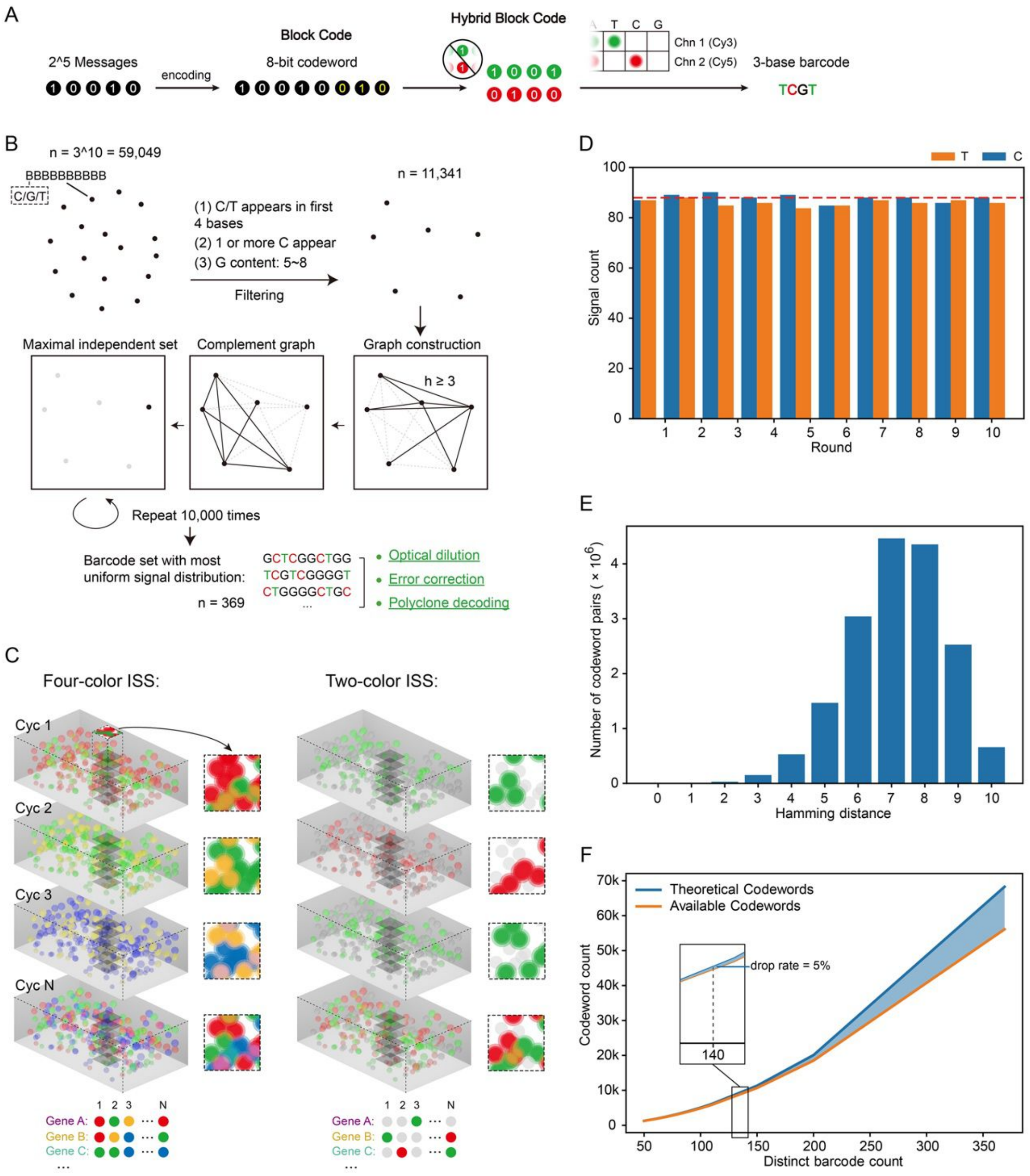


**Fig. S9. Probe design. (A)** Hybrid block code. n-bit block binary code was used to encode 2k genes (n > k), after which the block code was folded into n/2 double-binary code to shorten the read length. Dual "1" channels were avoided. To convert the final codeword into DNA barcode, two optical channels were used to represent two binary codeword channels (Chn) in which T and C were used to represent 1 in these two channels and G to represented 0. (**B)** Barcode design. All 59,049 barcode candidates were generated and filtered by 3 restrictions (in parenthesized enumerated list). The remaining 11,341 candidates were used as nodes for graph construction. Two nodes in the graph are connected by an edge if the Hamming distance between them is more than 2. After 10,000 trials, the maximal independent set (MIS) of the complement graph was determined. The final barcode list was selected from the MIS with the most uniform signal distribution. **(C)** Signal optical dilution from using label-free G bases in two-color in-situ sequencing (ISS) compared with typical four-color sequencing. **(D)** Barcode library signal counts. The horizontal dashed line indicates the mean signal count per channel in one reaction round. “C” and “T” are evenly distributed in each round in our 369-barcode library. **(E)** The Hamming distance distribution across the codewords. From the total 6,105 barcodes, 234 overlapping barcodes were eliminated (110 original barcodes + $\binom{110}{2}$ composite barcodes). Of all 17,231,385 pairs, 34,237 pairs (~0.2% drop rate) had a Hamming distance no greater than 2. The Hamming distances were calculated by 10-letter words constructed with the alphabet A, T, C, and G. **(F)** Drop rate of overlapping barcodes. The difference between theoretical and available codewords reflects the dropped barcodes (original barcodes + composite barcodes). For an acceptable 5% drop rate, the maximal subset size is 140 out of 369 original barcodes. Subsets sizes of 50, 60, 70, 80, 90, 100, 110, 150, 200, and 369 were used for this calculation.

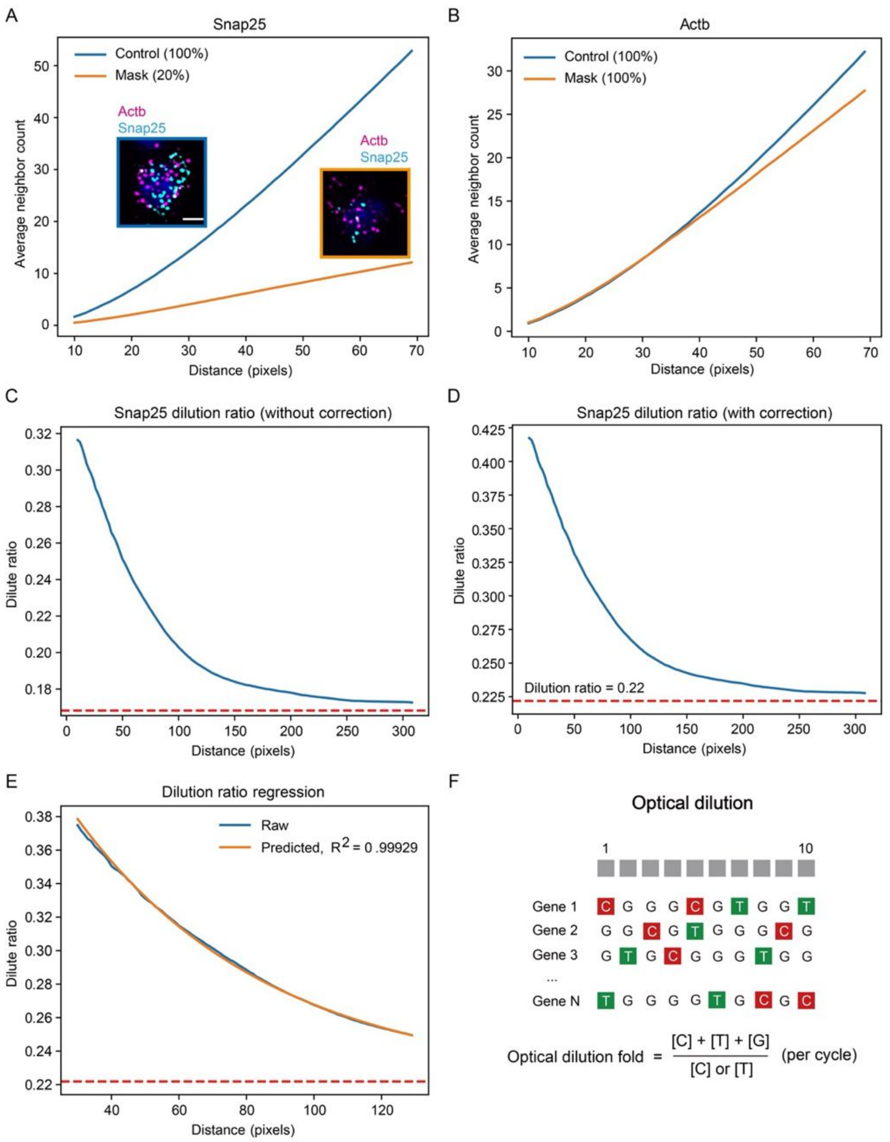


**Fig. S10. Signal dilution. (A-B)** Average neighbor counts of signals along a distance. Snap25 was masked by 80% in the mask group, while Actb was not masked in either group. Thus, the signal density of Actb can be used to correct an experimental difference. Pixel size is 0.1625 x 0.1625 μm$^2$. Scale bar: 5 μm **(C-D)** Dilution fold characterized by selective amplification with and without correction. **(E)** Curve fitting of dilution fold by selective amplification along a distance. **(F)** Optical dilution fold. The optical dilution fold produced by silent bits can be calculated by the total signal number (“C”+“T”+“G”) divided by the number of C or T signals in each cycle in the library.

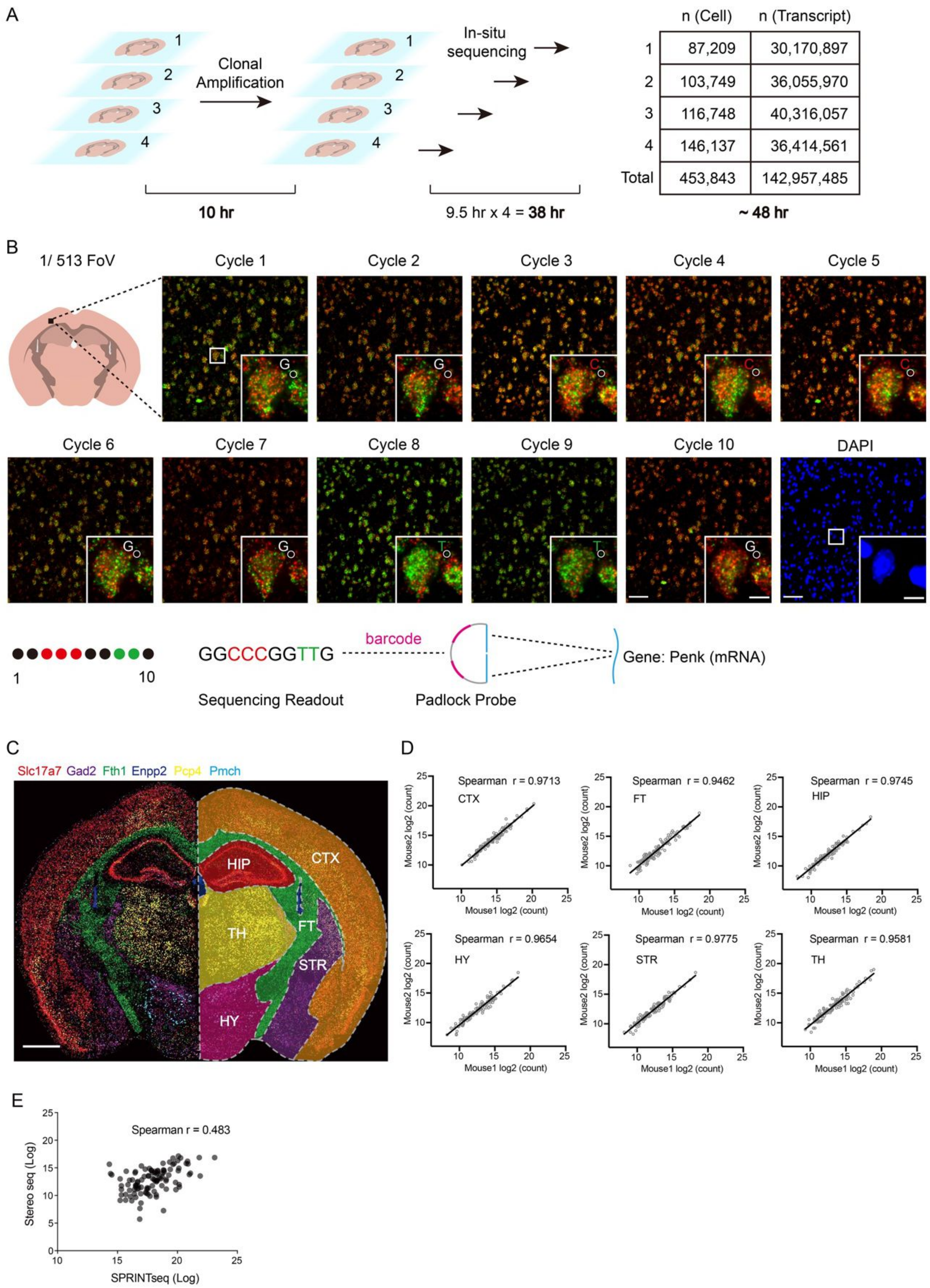


**Fig. S11. Raw sequencing results of a normal mouse brain slice. (A)** As stated in section 5.1, four slices of mouse whole brain were profiled in 48 hours. Multiple slices can be prepared for sequencing at the same time. The samples were sequentially mounted to our sequencing device for in-situ sequencing, and each take ~9.5 hours with a 40 x objective lens. This step can be finished as fast as 5 hours with a 20 x objective lens. **(B)** Raw image examples of mouse brain under 2 channels from 10 sequencing cycles followed by 1 DAPI cycle. The circled spot is an example of the mapping sequencing data-to-gene barcode. Scale bar: 10 μm. **(C)** Brain regions were identified by the distributions of 6 genes (Slc17a7, Gad2, Fth1, Enpp2, Pcp4, and Pmch). CTX: cerebral cortex, FT: fiber tracts, HIP: hippocampal region, HY: hypothalamus, STR: striatum, TH: thalamus. Density distribution (downsampled) of respective genes is shown with different color on the left, manually divided brain regions according to those 6 genes were shown on the right. Scale bar: 1mm. **(D)** Expression replicability of the different brain regions in 2 representative mouse brains. **(E)** Benchmark analysis through comparison with Stereo-seq. Both samples are coronal slice of mouse brain, the spots represent shared 104 genes.

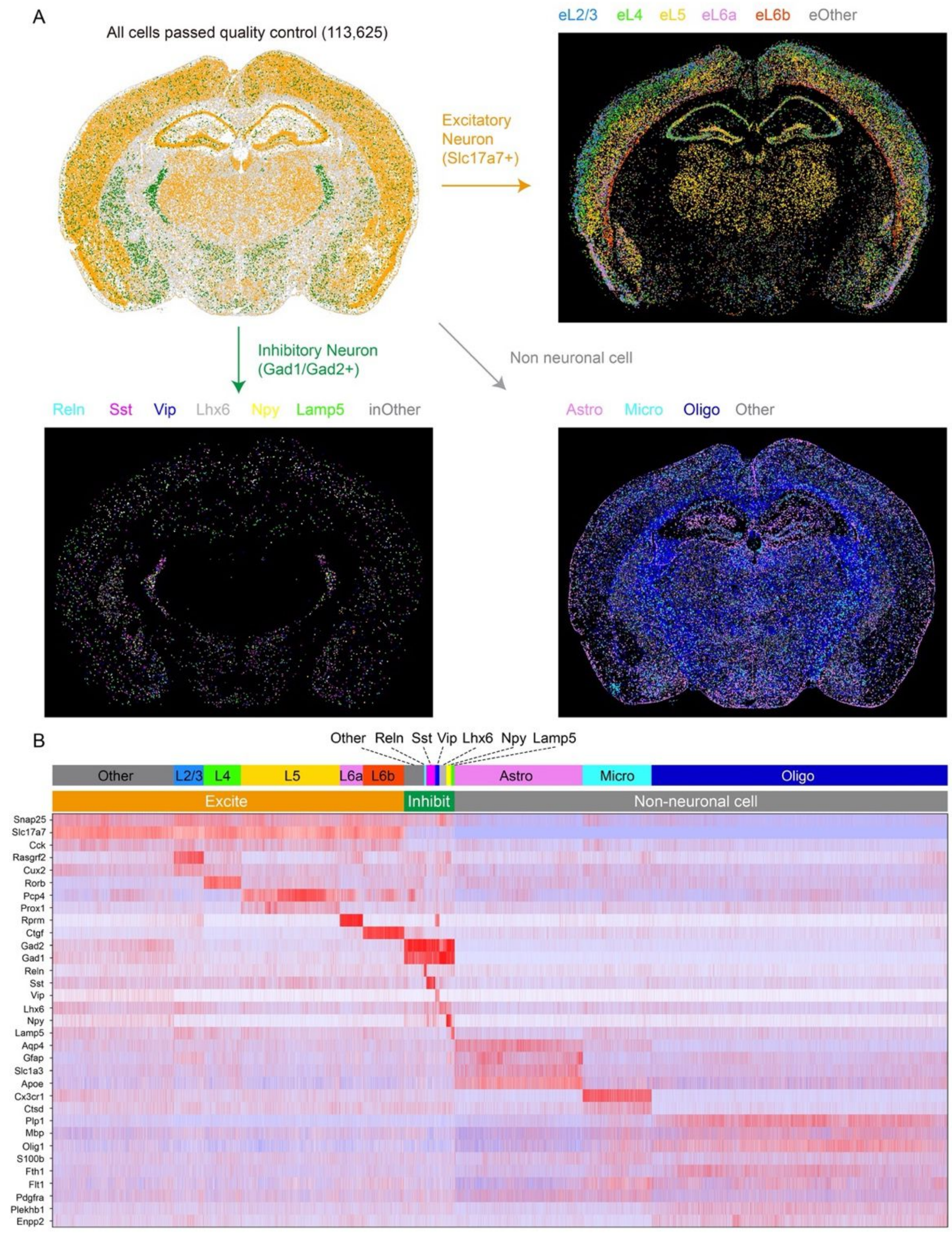


**Fig. S12. Cell classification of a brain coronal slice. (A)** Cell classification and projection. All cells that passed quality control were divided into three general categories (excitatory neurons, inhibitory neurons, and non-neuronal cells) based on marker gene expression, then the cells within each category were clustered. The colors of the cell types above each photo correspond to the dye colors in the photo. **(B)** Expression heatmap of all 17 cell types.

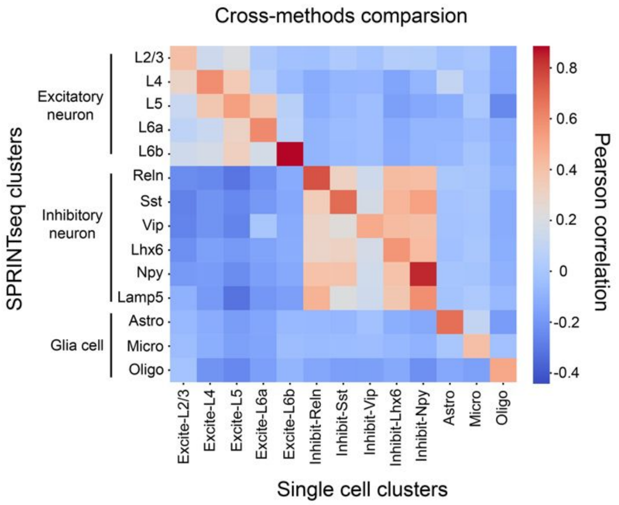


**Fig. S13. Clusters expression correlation between SPRINTseq's data and single cell RNA sequencing data.** The cell clusters from SPRINTseq were found in single cell data (combined by cortex, thalamus, hippocampus and hypothalamus data), and pearson correlation was calculated in each corresponding clusters according to the expression pattern of shared 104 gene.

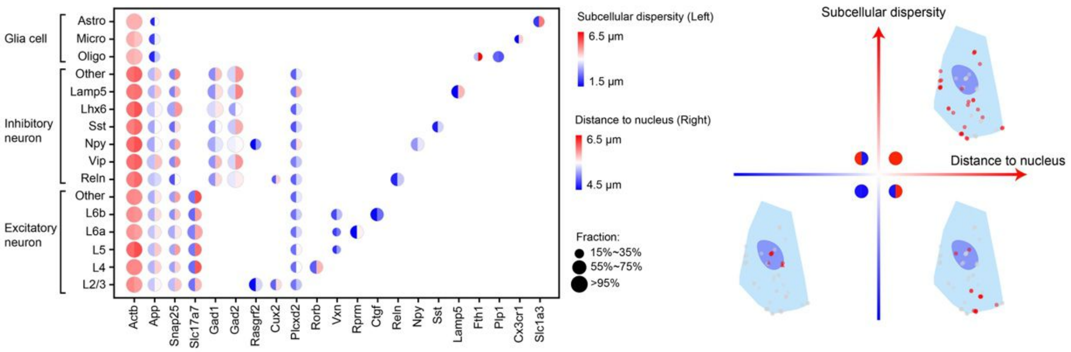


**Fig. S14. Subcellular RNA distributions.** Subcellular dispersity was calculated by the average distance to its centroid for each RNA species, and distance to the nucleus was calculated as the average RNA spot distance to a nucleus centroid for each RNA species. The circle sizes indicate expression proportions, and genes with an expression proportion less than 30% are not shown in the table. The coordinate system on the right shows a typical subcellular RNA distribution.

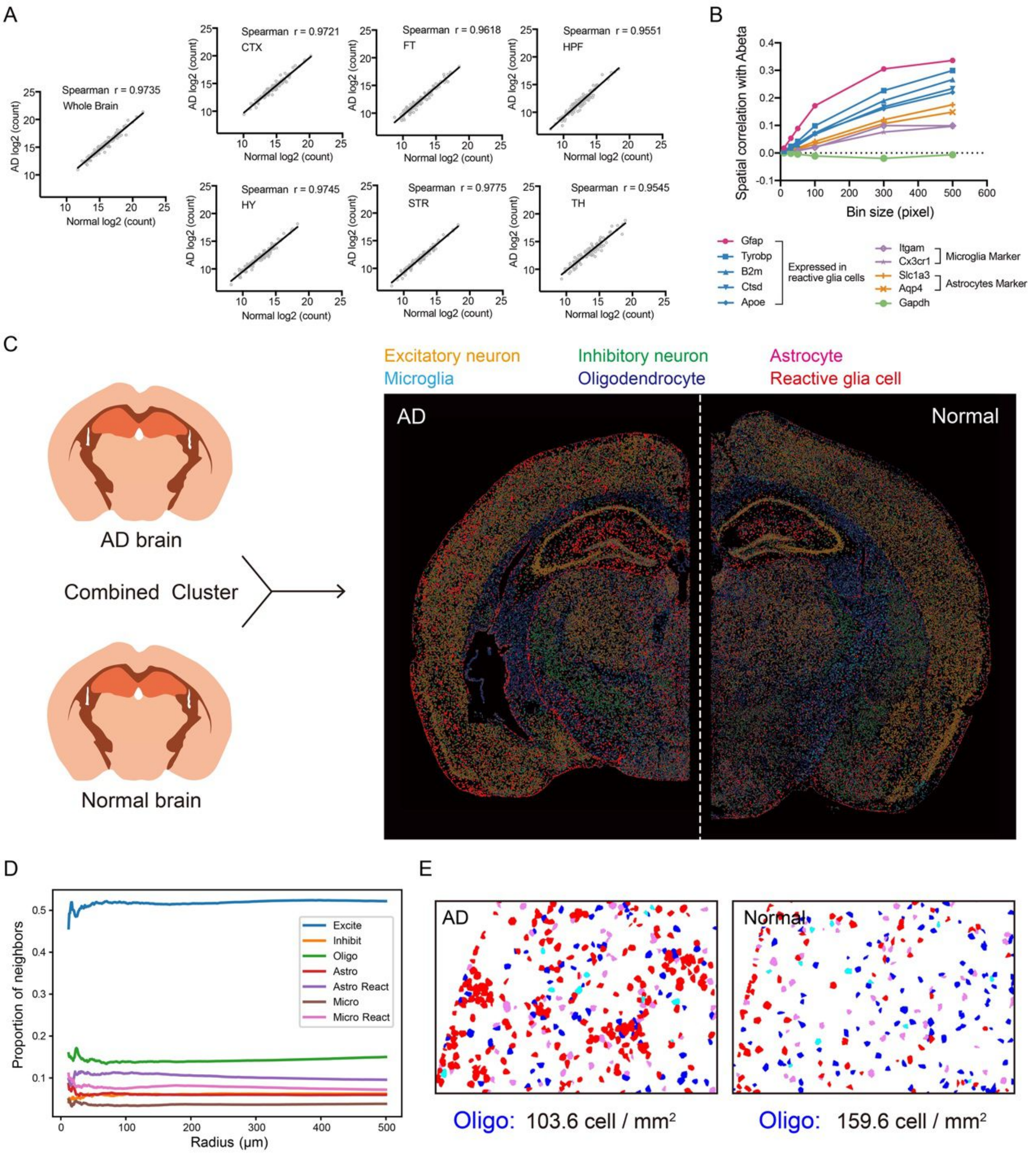


**Fig. S15. SPRINTseq analysis of mouse brain with Alzheimer's disease (AD). (A)** Spearman correlations between AD and normal mouse brain at the whole brain and brain region scales (See Fig. 10B for brain region names). **(B)** Spatial correlation analysis of specific genes and amyloid plaque (Amyloid β) within different bin sizes ($10 \times 10$, $30 \times 30$, $50 \times 50$, $100 \times 100$, $300 \times 300$, $500 \times 500$ px$^2$, where $1\text{px} = 0.1625\ \mu\text{m}$). The ranking of enriched genes did not change with bin size. Correlations between the housekeeping (control) gene (e.g., Gapdh) with amyloid plaque was about 0 (horizontal dotted line) in all bin sizes. In the whole gene panel, no significantly negative (smaller than -0.1) correlated genes were enriched. **(C)** Combined classification of AD and normal half mouse brains. Cells were combined together for type identification before being projected to its source brain. **(D)** Relative cell density changes as a function of distance to amyloid plaque, but with a random jitter (range: 0~500 μm) applied to plaque coordinates. The trends of neuron and reactive glia cells observed without jitter disappeared, suggesting an authentic correlation between the cellular distribution and location of amyloid plaques. **(E)** Oligodendrocyte density (red dots) decreased by ~35% in the AD mouse brain cortex compared to that in the normal mouse brain.

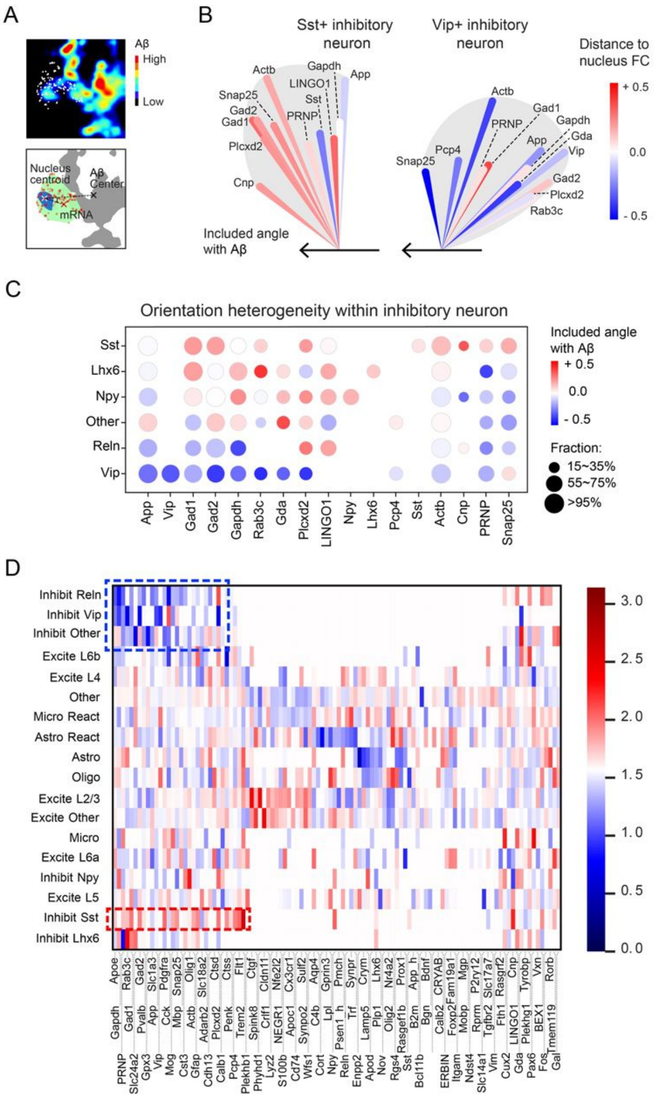


**Fig. S16. SPRINTseq analysis of mouse brain with Alzheimer's disease (AD).** **(A)** Diagram of mRNA orientation characterization. In a given cell, orientation is calculated as the included angle formed by amyloid plaque, the nucleus centroid, and the mRNA. This value is calculated only within cells that are close to amyloid plaque. A smaller included angle might suggest a trend for mRNA to approach plaque. mRNA locations are white in the top image and red in the bottom diagram. **(B)** A combination of mRNA orientation and the change of distance to the nucleus centroid. The change of distance to a nucleus centroid was calculated as a subcellular dispersion change. This change is also calculated by the value of cells close to amyloid plaque divided by cells far from amyloid plaque. Major expressed genes in two cell types were chosen as examples. The length of the scalloped area represents the expression level of each gene (log2). **(C)** Heatmap of mRNA orientation in inhibitory neurons. Y axis shows different subtypes within inhibitory neuron. The genes listed are major expressed genes in inhibitory neurons. **(D)** The subcellular mRNA polarity pattern in cells that are within 25 μm to Amyloid β. Y axis shows different cell types. The blue and red dashed boxes enclose results that display great subcellular mRNA distribution heterogeneity in inhibitory cells.

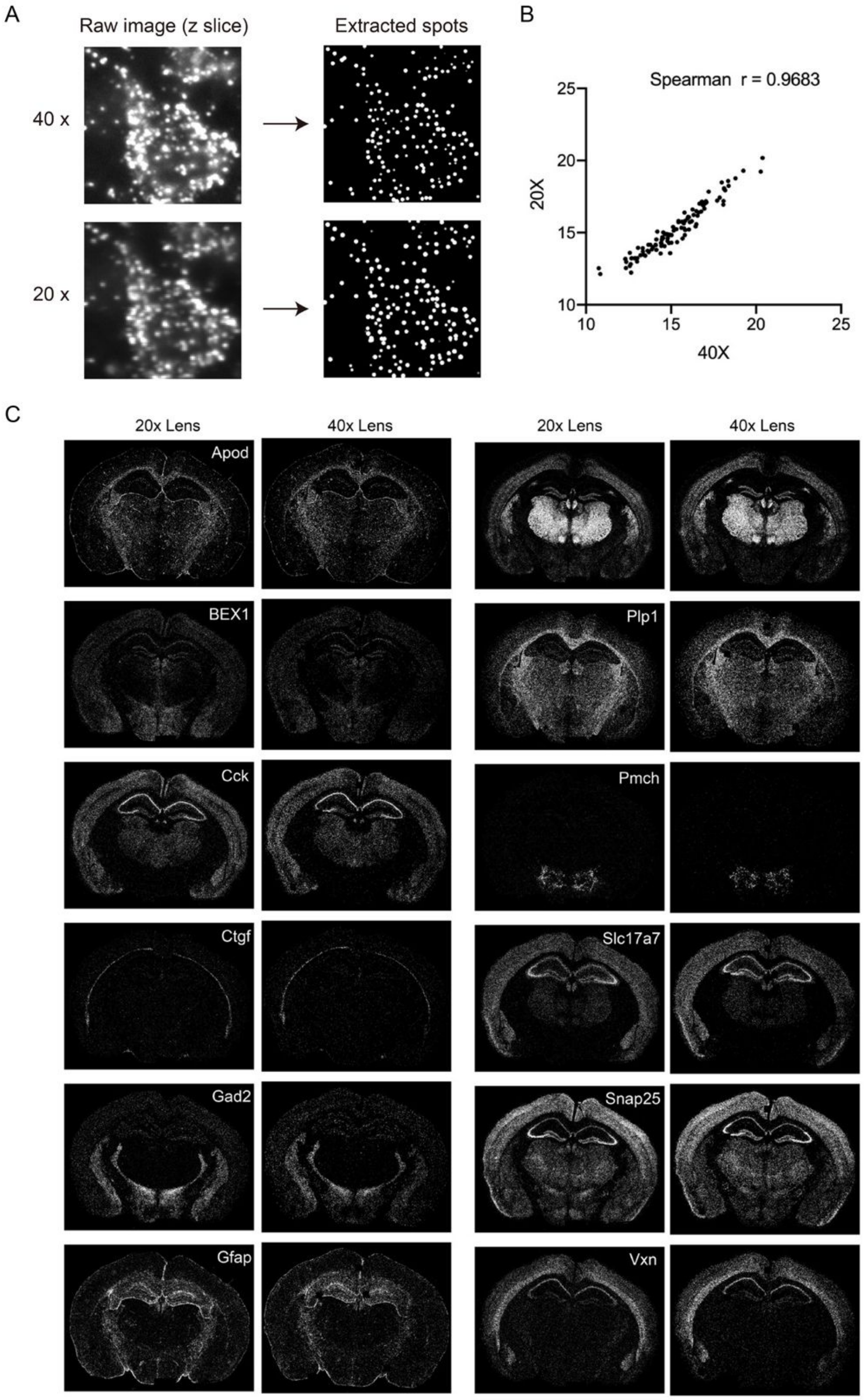


**Fig. S17. Improving SPRINTseq throughput by using low magnification lens (20x). (A)** Direct comparison of image quality taken by 20x and 40x lens. **(B)** Spearman correlations of gene expression levels between results imaged by 20x and 40x lens respectively. **(C)** Sequencing results imaged by 20x lens with those by 40x lens, genes example with specific distribution patterns are listed.

### Extra Tables

### Extra Table 1. The sequences of Padlock probes

| Gene | Padlock Probe Sequence |
|---|---|
| Slc17a7 | GGACGTAAAGAAGCGCCTCCTCCCTACACGACGCTCTTCCGATCTGCTCGGCTGGCATTCCTGCTGAACCGCTCTTCCGAGCTCGGCTGGATGATGGCATAGACGGGCAT |
| Gad1 | TCAGCTACTGACAGAGCTGTTCCCTACACGACGCTCTTCCGATCTGCTGGTGCGCCATTCCTGCTGAACCGCTCTTCCGAGCTGGTGCGCAGTGAGATGGCCTAGATGTG |
| Gad2 | TACAGTCATACTGAGGATCATCCCTACACGACGCTCTTCCGATCTTGGCCGGCGTCATTCCTGCTGAACCGCTCTTCCGATGGCCGGCGTAGACAGCAAGGGCCCAAAGC |
| Lhx6 | GTGGGTAGAGCTGGCTCTGTTCCCTACACGACGCTCTTCCGATCTTGCGGGCGGCCATTCCTGCTGAACCGCTCTTCCGATGCGGGCGGCGTTGGCCTCTGCTCTTAGGA |
| Adarb2 | CACTAGAGAGGCCCTTGGAATCCCTACACGACGCTCTTCCGATCTCCTGGGGTGCCATTCCTGCTGAACCGCTCTTCCGACCTGGGGTGCCTGGTGAGACTTGATCACAC |
| Lamp5 | AGCACCCAGGATGCAGGAGCTCCCTACACGACGCTCTTCCGATCTCGTGCCCGGGCATTCCTGCTGAACCGCTCTTCCGACGTGCCCGGGCCGCCGCACACTGCAGAGGA |
| Vip | CTTGACACATCATAATAGGGTCCCTACACGACGCTCTTCCGATCTGGTCGGTCGTCATTCCTGCTGAACCGCTCTTCCGAGGTCGGTCGTCATCAGCATGCCTGGCATTT |
| Pax6 | GCCAAGAGCTGGAGGTGGGATCCCTACACGACGCTCTTCCGATCTTGCTGTGGTGCATTCCTGCTGAACCGCTCTTCCGATGCTGTGGTGGCCCTTGGTTAAAGTCTTCT |
| Pvalb | TCCTTTGACTTATCTCACTTTCCCTACACGACGCTCTTCCGATCTGGCGCGGTGCCATTCCTGCTGAACCGCTCTTCCGAGGCGCGGTGCCAGGAGATATCGGGGCGTTG |
| Sst | GCCTGGGGCAAATCCTCGGGTCCCTACACGACGCTCTTCCGATCTGTTCGGGGCGCATTCCTGCTGAACCGCTCTTCCGAGTTCGGGGCGTCATCTCGTCCTGCTCAGCT |
| Npy | GGTGGAAACTTGGAAAAGTCTCCCTACACGACGCTCTTCCGATCTTGCCTGGGGTCATTCCTGCTGAACCGCTCTTCCGATGCCTGGGGTAGGGGATGAGATGAGATGAG |
| Cux2 | TTTGGTTGGCTGAGACCTCCTCCCTACACGACGCTCTTCCGATCTTGGGGCCCGCCATTCCTGCTGAACCGCTCTTCCGATGGGGCCCGCCTGCCGCTCCAGGTCAGCGA |
| Plcxd2 | GCCTCCTCCATGGTGTCATATCCCTACACGACGCTCTTCCGATCTCCTGGCGCGGCATTCCTGCTGAACCGCTCTTCCGACCTGGCGCGGTTCCCACGCATCCTGTGTCA |
| Pcp4 | GTATTGAGTGAGGATGACTTTCCCTACACGACGCTCTTCCGATCTCGCTCTGGGGCATTCCTGCTGAACCGCTCTTCCGACGCTCTGGGGTCAGGTTTGTAGCAGGGTGT |
| Ctgf | GGCTTTACGCCATGTCTCCGTCCCTACACGACGCTCTTCCGATCTGCTGGCTTGGCATTCCTGCTGAACCGCTCTTCCGAGCTGGCTTGGGTTCGTGTCCCTTACTTCCT |
| Sulf2 | CAAGGATGTAGCACCGATGGTCCCTACACGACGCTCTTCCGATCTCGTGGGTCCGCATTCCTGCTGAACCGCTCTTCCGACGTGGGTCCGGCACTGGACTGTGTCATTCT |
| Rorb | TACAGATGTGAGGTCATAGATCCCTACACGACGCTCTTCCGATCTGGTGTGGCCCCATTCCTGCTGAACCGCTCTTCCGAGGTGTGGCCCCTATAGGTAAACAAGTTGGG |
| Ndst4 | GGTCTATGAGGATAGTAATGTCCCTACACGACGCTCTTCCGATCTGCGCTCGGTGCATTCCTGCTGAACCGCTCTTCCGAGCGCTCGGTGAGAATATGCCCTGTCCGAGG |
| Nov | GTGGGACAACTTCATTATGTTCCCTACACGACGCTCTTCCGATCTCTGGGGCTGCCATTCCTGCTGAACCGCTCTTCCGACTGGGGCTGCATATGATCACAGCAACATAC |
| Slc1a3 | AGATGGATACATTGTAGAACTCCCTACACGACGCTCTTCCGATCTTCGTCGGGGTCATTCCTGCTGAACCGCTCTTCCGATCGTCGGGGTCTGGGAAAGTGAGCCCAGGG |
| Mgp | CAGCGTTGTAGCCGTAGACCTCCCTACACGACGCTCTTCCGATCTGGTGCTCGGCCATTCCTGCTGAACCGCTCTTCCGAGGTGCTCGGCCCTGAAGTAGCGGTTGTAGG |
| Aqp4 | TCAGGACAGAAGACATACTCTCCCTACACGACGCTCTTCCGATCTGGGCCGCTCGCATTCCTGCTGAACCGCTCTTCCGAGGGCCGCTCGGGCGACGTTTGAGCTCCACA |

| | |
|---|---|
| **Actb** | CATGATGGAATTGAATGTAGTCCCTACACGACGCTCTTCCGATCTCGCCGGGGGGCATTC<br>CTGCTGAACCGCTCTTCCGACGCCGGGGGGCGGATGTCAACGTCACACTT |
| **Gapdh** | TTGATGTTAGTGGGGTCTCGTCCCTACACGACGCTCTTCCGATCTTGCGGGTCGGCATTC<br>CTGCTGAACCGCTCTTCCGATGCGGGTCGGCAGCACCGGCCTCACCCCAT |
| **Snap25** | ACAAAGCCCGCAGAATTTTCTCCCTACACGACGCTCTTCCGATCTTTGGCGCCGGCATTC<br>CTGCTGAACCGCTCTTCCGATTGGCGCCGGAGCTTGTTACAGGGACACAC |
| **Foxp2** | TTGCAGCTGTAGCCTTTGACTCCCTACACGACGCTCTTCCGATCTTGGGGCGCCGCATTC<br>CTGCTGAACCGCTCTTCCGATGGGGCGCCGTGGTTAGAGTTGGCTTTGGT |
| **Pdgfra** | AGGCTGGCCGATCGTACAGCTCCCTACACGACGCTCTTCCGATCTGTCGCTGGGCCATTC<br>CTGCTGAACCGCTCTTCCGAGTCGCTGGGCCATGGATTTCTTCTTGTAGG |
| **Mog** | GGAGAACGGTACCAACCCACTCCCTACACGACGCTCTTCCGATCTGCGCGGGGGCCATTC<br>CTGCTGAACCGCTCTTCCGAGCGCGGGGGCGGTGAACCACTCTTGAGAAG |
| **Itgam** | CTCACTGAGTCGTCCACGCATCCCTACACGACGCTCTTCCGATCTCGGGGTTTTGCATTC<br>CTGCTGAACCGCTCTTCCGACGGGGTTTTGTGAGGCGCAGGATGATGGGG |
| **Flt1** | GAGCAGGTCAGTGCATGCAGTCCCTACACGACGCTCTTCCGATCTGCGGGCGGCGCATTC<br>CTGCTGAACCGCTCTTCCGAGCGGGCGGCGTTTCCTCCATAAGAGAGACA |
| **Bgn** | TAGCTGGGATTGACAGGTAGTCCCTACACGACGCTCTTCCGATCTTTCTGGTGGGCATTC<br>CTGCTGAACCGCTCTTCCGATTCTGGTGGGGAGTTAGTACAGCCAGGTTG |
| **Cdh13** | CATTCACTGTCAGCAGCACATCCCTACACGACGCTCTTCCGATCTCGGGTGCGCCCATTC<br>CTGCTGAACCGCTCTTCCGACGGGTGCGCCCAGGGAGTCTGGGTCTGTGG |
| **Ctsd** | CAGCTCCTTCACCTCTTCCATCCCTACACGACGCTCTTCCGATCTGCCGGGTCTGCATTC<br>CTGCTGAACCGCTCTTCCGAGCCGGGTCTGACTGCCCCGATGGCCTTCTG |
| **Cst3** | TCCTCATTGGCATCTGCCTCTCCCTACACGACGCTCTTCCGATCTGTGGGCGCGGCATTC<br>CTGCTGAACCGCTCTTCCGAGTGGGCGCGGCCAACGCTCGCCGCACGCCT |
| **Apoe** | TGCTGGGTCTGTTCCTCCATTCCCTACACGACGCTCTTCCGATCTCCGGGGCGCGCATTC<br>CTGCTGAACCGCTCTTCCGACCGGGGCGCGTCTCCGCCTGCAGGCGTATT |
| **Gfap** | CTGGAGGTTGGAGAAAGTCTTCCCTACACGACGCTCTTCCGATCTTGCCCGGGCGCATTC<br>CTGCTGAACCGCTCTTCCGATGCCCGGGCGTCCAGGCTGGTTTCTCGGAT |
| **Plp1** | GCAAACACCAGGAGCCATACTCCCTACACGACGCTCTTCCGATCTTGCCGCGTGGCATTC<br>CTGCTGAACCGCTCTTCCGATGCCGCGTGGACACAGGTACAGCCGAGCAG |
| **Mbp** | CCCTTGGGATGGAGGTGGTGTCCCTACACGACGCTCTTCCGATCTGCGTCCGGGGCATTC<br>CTGCTGAACCGCTCTTCCGAGCGTCCGGGGCGGCTGTCTCTTCCTCCCTT |
| **Cnp** | GACGTGAGCTCGGCTCCCTGTCCCTACACGACGCTCTTCCGATCTGCTCTGGGGTCATTC<br>CTGCTGAACCGCTCTTCCGAGCTCTGGGGTTCGGCCGCACAGCCTAGGGT |
| **App** | TGTCTCTACAAGCTGCTGTCTCCCTACACGACGCTCTTCCGATCTCGCTGGGCGCCATTC<br>CTGCTGAACCGCTCTTCCGACGCTGGGCGCGCTTCAACTCTGGCCATGTG |
| **Olig2** | GCGGCTGTTGATCTTCAGGCTCCCTACACGACGCTCTTCCGATCTGGGCGTGCCCCATTC<br>CTGCTGAACCGCTCTTCCGAGGGCGTGCCCTCGTGCATGCGCTTGCGTTC |
| **Tyrobp** | ACAGTCGCATCTTGGGAAAGTCCCTACACGACGCTCTTCCGATCTTGGTGTCGGGCATTC<br>CTGCTGAACCGCTCTTCCGATGGTGTCGGGACACCAGGGCTCACGGAAGA |
| **Lpl** | GTCTCAAATGAAATACAAAGTCCCTACACGACGCTCTTCCGATCTGGCGCCTTGGCATTC<br>CTGCTGAACCGCTCTTCCGAGGCGCCTTGGCAGAGGCATGGCGGAGATGA |
| **Trem2** | CATGCAGGCTGGATTGACTCTCCCTACACGACGCTCTTCCGATCTGCCGGTCGGGCATTC<br>CTGCTGAACCGCTCTTCCGAGCCGGTCGGGCTTGGTGGAGGAGGGGAGAG |
| **Psen1_human** | TTTGTTATAGTCAAAGAAGATCCCTACACGACGCTCTTCCGATCTGGGCCTCCGGCATTC<br>CTGCTGAACCGCTCTTCCGAGGGCCTCCGGCACCTTTGTCCTCCCCAGAT |
| **App_human** | GATCCGCCGCGTCCTTGCTCTCCCTACACGACGCTCTTCCGATCTTGGGCCCTGGCATTC<br>CTGCTGAACCGCTCTTCCGATGGGCCCTGGGTGCGCTGCTGTGCGAGTGG |

| | |
|---|---|
| **Reln** | ATCTTGTAGCGTTGGTTTGGTCCCTACACGACGCTCTTCCGATCTGGGCGCTGTCCATTC<br>CTGCTGAACCGCTCTTCCGAGGGCGCTGTCGTACGGTTGCCAGAGTCTGA |
| **Calb1** | GCTTCATCGAAACCGAGGAATCCCTACACGACGCTCTTCCGATCTTGTGTGTGGCCATTC<br>CTGCTGAACCGCTCTTCCGATGTGTGTGGCGTATGATACTGACCACAGCG |
| **Slc14a1** | AGCAGCGTGCAGGCACATGATCCCTACACGACGCTCTTCCGATCTTTGGGGTGGCCATTC<br>CTGCTGAACCGCTCTTCCGATTGGGGTGGCACACCCAGCAACGATCCAAT |
| **Vxn** | CGGTGAAAGCTCCAGCAGGGTCCCTACACGACGCTCTTCCGATCTCTGTGCGTGGCATTC<br>CTGCTGAACCGCTCTTCCGACTGTGCGTGGTCCTCCAGTGGCTAGGATCT |
| **Calb2** | GCCCACGTGCTGCCTGAAGCTCCCTACACGACGCTCTTCCGATCTGGCGGGCTCGCATTC<br>CTGCTGAACCGCTCTTCCGAGGCGGGCTCGTCCATAAACTCAGCGCTGGA |
| **Penk** | CGGAGGAGTTGGCCAAGGTGTCCCTACACGACGCTCTTCCGATCTGGCCCGGTTGCATTC<br>CTGCTGAACCGCTCTTCCGAGGCCCGGTTGCAGTAGCTCTTTCAGCAGAT |
| **Rgs4** | GCACACCCTGAGCACCCAGGTCCCTACACGACGCTCTTCCGATCTCGCGGTGTGGCATTC<br>CTGCTGAACCGCTCTTCCGACGCGGTGTGGACCAACCCAGCTGCTAATCA |
| **Rprm** | TCCGTGATGGTGAGGGGTGCTCCCTACACGACGCTCTTCCGATCTTCTGGGGTCGCATTC<br>CTGCTGAACCGCTCTTCCGATCTGGGGTCGGACAGGTTTGCGTTGCTCGG |
| **Slc24a2** | CGTAAAGAAACGATGGAACATCCCTACACGACGCTCTTCCGATCTGTGCTGTGCGCATTC<br>CTGCTGAACCGCTCTTCCGAGTGCTGTGCGGAGTGACTGCTTCCAAGTAG |
| **Wfs1** | AGATCACCGTGGTGACCATGTCCCTACACGACGCTCTTCCGATCTGTCGTGGCGCCATTC<br>CTGCTGAACCGCTCTTCCGAGTCGTGGCGCGAAAAGCAGGGGTACGCCGC |
| **Crym** | ACGGTGCGCACTGGCTGCATTCCCTACACGACGCTCTTCCGATCTGGGCGCGCTTCATTC<br>CTGCTGAACCGCTCTTCCGAGGGCGCGCTTGGTGCTTGGCTACAGGCACC |
| **Gda** | CCTCAAGCACGTTCAGTAAGTCCCTACACGACGCTCTTCCGATCTGGGTCCCGCGCATTC<br>CTGCTGAACCGCTCTTCCGAGGGTCCCGCGCTTCACTTTATGCTTCAGGA |
| **Bcl11b** | CCAACAGCTCCGGCATCACCTCCCTACACGACGCTCTTCCGATCTGTGTGGGGTCCATTC<br>CTGCTGAACCGCTCTTCCGAGTGTGGGGTCGTGCACCCGCGTCCCCACAG |
| **Cck** | TCCCGGTCACTTATTCTATGTCCCTACACGACGCTCTTCCGATCTCTTGGTTGGGCATTC<br>CTGCTGAACCGCTCTTCCGACTTGGTTGGGAATCCATCCAGCCCATGTAG |
| **Fam19a1** | CCGTTCTAACCTCTTCCAGGTCCCTACACGACGCTCTTCCGATCTGTGCGCTTGGCATTC<br>CTGCTGAACCGCTCTTCCGAGTGCGCTTGGATTCGATTGATATATAATAA |
| **Rab3c** | ACAAATGGTGCACAGCAAAATCCCTACACGACGCTCTTCCGATCTGTGTGCTGTGCATTC<br>CTGCTGAACCGCTCTTCCGAGTGTGCTGTGCTTTTAAGTAAGACGTGGTC |
| **Synpr** | ATAAAGTCAATGAGTGGGCCTCCCTACACGACGCTCTTCCGATCTGTGTTGTCGGCATTC<br>CTGCTGAACCGCTCTTCCGAGTGTTGTCGGATGAAAAGACTACAGTGACG |
| **Cort** | GTAGCGAGCATTACTGAAACTCCCTACACGACGCTCTTCCGATCTGCGGGTGCTGCATTC<br>CTGCTGAACCGCTCTTCCGAGCGGGTGCTGCCTTAGTTGACCTGTTGTCG |
| **Enpp2** | CTGCCCCACGGTCTTGTCAATCCCTACACGACGCTCTTCCGATCTTCGTGGGGCGCATTC<br>CTGCTGAACCGCTCTTCCGATCGTGGGGCGTGTTTCAGTCCGTCCATTAA |
| **Rasgrf2** | TCAACCGCTCCAAGAGGCGCTCCCTACACGACGCTCTTCCGATCTTCTTTGGGGGCATTC<br>CTGCTGAACCGCTCTTCCGATCTTTGGGGGGCTCAGAAACCGCAAGTCCG |
| **Nr4a2** | GCGTTGTCACCGCAAACAGCTCCCTACACGACGCTCTTCCGATCTCTGGTGGGGTCATTC<br>CTGCTGAACCGCTCTTCCGACTGGTGGGGTCACCGTAGTGCTGACAGGCC |
| **Fos** | ACCTGCCTGCAAGATCCCCGTCCCTACACGACGCTCTTCCGATCTGTTGGCGGGCCATTC<br>CTGCTGAACCGCTCTTCCGAGTTGGCGGGCTTTATTTTGGCAGCCCACCG |
| **Lyz2** | GCCATGCCACCCATGCTCGATCCCTACACGACGCTCTTCCGATCTGTCGGTTCGGCATTC<br>CTGCTGAACCGCTCTTCCGAGTCGGTTCGGTCGGTTTTGACAGTGTGCTC |
| **B2m** | CACGCCACCCACCGGAGAATTCCCTACACGACGCTCTTCCGATCTGCGGTCGCGCCATTC<br>CTGCTGAACCGCTCTTCCGAGCGGTCGCGCCCCTCAAATTCAAGTATACT |

| | |
|---|---|
| **Mobp** | TCCTCGCTCTGGTCTGCCTCTCCCTACACGACGCTCTTCCGATCTGTGGCGTGTTCATTC<br>CTGCTGAACCGCTCTTCCGAGTGGCGTGTTCGCGGGGCTGAGTTTCCGTC |
| **Cldn11** | CGAGCCTGGAGTGGCCAAGTTCCCTACACGACGCTCTTCCGATCTGTCGTGCGTGCATTC<br>CTGCTGAACCGCTCTTCCGAGTCGTGCGTGCCCTGCATCCGAATGGGCCA |
| **Apod** | CGCTTGATCTGTTGTTGTCATCCCTACACGACGCTCTTCCGATCTGTTCTGGCGGCATTC<br>CTGCTGAACCGCTCTTCCGAGTTCTGGCGGTACAGGAAGTCCGGGCAGTT |
| **Trf** | CCCTCCGGGCACACGCCTTCTCCCTACACGACGCTCTTCCGATCTCTTTGGGGGGCATTC<br>CTGCTGAACCGCTCTTCCGACTTTGGGGGGCTGGCGAGTTGTCGATCGAG |
| **Fth1** | CCCAGAGTCGCCGCGGTTTCTCCCTACACGACGCTCTTCCGATCTGGCGTCGCGTCATTC<br>CTGCTGAACCGCTCTTCCGAGGCGTCGCGTTCCTGCGCCAGACGTTCTCG |
| **Plekhb1** | CCTGATACGGGCTGTAGACGTCCCTACACGACGCTCTTCCGATCTGCGGGGGCCTCATTC<br>CTGCTGAACCGCTCTTCCGAGCGGGGGCCTTGGCACCACCTCATAGTAGT |
| **LING01** | GCAGGTGAAGTAGTTGGGTATCCCTACACGACGCTCTTCCGATCTGCCCGGCGCGCATTC<br>CTGCTGAACCGCTCTTCCGAGCCCGGCGCGTCCCGGATGTGGGCCCGGCG |
| **CRYAB** | CCTTCGGCCACCCTCCTTCCTCCCTACACGACGCTCTTCCGATCTGTGGCGGTCTCATTC<br>CTGCTGAACCGCTCTTCCGAGTGGCGGTCTACTTCCCTGAGCCCCTTCTA |
| **PRNP** | AGCAACCAGAACAACTTCGTTCCCTACACGACGCTCTTCCGATCTGCGGGGGTTCCATTC<br>CTGCTGAACCGCTCTTCCGAGCGGGGGTTCACAGGCCAGTGGATCAGTAC |
| **CNTNAP2** | AGTGGAGATCGTGGTGAGGTTCCCTACACGACGCTCTTCCGATCTGTGGGCTGCTCATTC<br>CTGCTGAACCGCTCTTCCGAGTGGGCTGCTTTTGATGTTGGAAACGGGCC |
| **ERBIN** | TAGCCCACATTTCTCCATGCTCCCTACACGACGCTCTTCCGATCTTCGGCCGCGGCATTC<br>CTGCTGAACCGCTCTTCCGATCGGCCGCGGTAGTCGGTCATTCGGGGAGA |
| **NEGR1** | GAGGGTGCAGGTGTGCCGCCTCCCTACACGACGCTCTTCCGATCTGTGCTTGGGTCATTC<br>CTGCTGAACCGCTCTTCCGAGTGCTTGGGTGCAGTGGACTGATACGATGC |
| **BEX1** | ATACGGATCTTCCCATGCAATCCCTACACGACGCTCTTCCGATCTGGTTGGTTGCCATTC<br>CTGCTGAACCGCTCTTCCGAGGTTGGTTGCTATTAGGTTCACAATAGGTA |
| **C4b** | CGGGGATAATAGCAGGCGAATCCCTACACGACGCTCTTCCGATCTGGCTTTCGGGCATTC<br>CTGCTGAACCGCTCTTCCGAGGCTTTCGGGCCGTGAAGCCATACTCCACT |
| **Prox1** | GAATAAGGTGAGATGTCGGATCCCTACACGACGCTCTTCCGATCTCGTCGCGGTGCATTC<br>CTGCTGAACCGCTCTTCCGACGTCGCGGTGCTTCCTGCATTGCGCTTCCT |
| **Crlf1** | CGTGGAGGACAGCGTGGACTTCCCTACACGACGCTCTTCCGATCTCGGGTCCGTGCATTC<br>CTGCTGAACCGCTCTTCCGACGGGTCCGTGAAGTACCAGATCCGCTACCG |
| **Spink8** | ATGTGGCAGCAACCAAGTGATCCCTACACGACGCTCTTCCGATCTGCGGTCCTGGCATTC<br>CTGCTGAACCGCTCTTCCGAGCGGTCCTGGTTCAAGGTCAGTGAGCCTAT |
| **Plekhg1** | CATTGGTGCCAGGATCGCTGTCCCTACACGACGCTCTTCCGATCTGGTCGTCGGTCATTC<br>CTGCTGAACCGCTCTTCCGAGGTCGTCGGTAAGAGGCAAGGAGGGCCCGC |
| **Synpo2** | GTACTTCCAACTGGCGGGCATCCCTACACGACGCTCTTCCGATCTGTCCGGGTGTCATTC<br>CTGCTGAACCGCTCTTCCGAGTCCGGGTGTGGTGCTCGGACGTTGGAGGA |
| **Pmch** | CAGAGAAGGGGCGACAACGGTCCCTACACGACGCTCTTCCGATCTTGGCCTGGTGCATTC<br>CTGCTGAACCGCTCTTCCGATGGCCTGGTGTCGTCGTTTTTGTATTGTTC |
| **Slc18a2** | CTCCATGATGCCTATCATGGTCCCTACACGACGCTCTTCCGATCTGGCGCTGGTGCATTC<br>CTGCTGAACCGCTCTTCCGAGGCGCTGGTGGCCATTGGGATGGTGGACTC |
| **Cd74** | CATGGATAACATGCTCCTTGTCCCTACACGACGCTCTTCCGATCTCTGGCTGCGGCATTC<br>CTGCTGAACCGCTCTTCCGACTGGCTGCGGTTGCTGATGCGTCCAATGTC |
| **Ctss** | ACGCCAGCCATTCCTCCTTCTCCCTACACGACGCTCTTCCGATCTCCGCCGGGCGCATTC<br>CTGCTGAACCGCTCTTCCGACCGCCGGGCGGCCTGTCTCTGTGGGCATCG |
| **Nfe2l2** | TGGCATCATCAGTGGAGAGGTCCCTACACGACGCTCTTCCGATCTGGGCGGCGCTCATTC<br>CTGCTGAACCGCTCTTCCGAGGGCGGCGCTGTCTAAGGAGGTCAGCTGGC |

| | |
|---|---|
| **Phyhd1** | TGCCTCTGGTTCATCCCTGGTCCCTACACGACGCTCTTCCGATCTTTCGGGGGGTCATTC<br>CTGCTGAACCGCTCTTCCGATTCGGGGGGTATGCCATGCTGGAGAACGGT |
| **S100b** | TTCTGCTAGGCTAGGCATTCTCCCTACACGACGCTCTTCCGATCTGGGTTCGGTCCATTC<br>CTGCTGAACCGCTCTTCCGAGGGTTCGGTCTTAAGTTAGCACATCATGAT |
| **Tgfbr2** | CCGTCAGGAACTGCAGGATGTCCCTACACGACGCTCTTCCGATCTTGGTTCTGGGCATTC<br>CTGCTGAACCGCTCTTCCGATGGTTCTGGGCTCTGTCTTCCGCTCCTCGG |
| **Vim** | CAACCTGGCCGAGGACATCATCCCTACACGACGCTCTTCCGATCTGGTCGCCGCGCATTC<br>CTGCTGAACCGCTCTTCCGAGGTCGCCGCGCGTGTCGAGGTGGAGCGGGA |
| **Gprin3** | CCAGAGACCTCCTTAGCCCCTCCCTACACGACGCTCTTCCGATCTCTGGGTGGTGCATTC<br>CTGCTGAACCGCTCTTCCGACTGGGTGGTGCAGCATCTTGCCAAGCCCTG |
| **Bdnf** | CTCATAGACATGTTTGCGGCTCCCTACACGACGCTCTTCCGATCTGGTGCGGTCGCATTC<br>CTGCTGAACCGCTCTTCCGAGGTGCGGTCGGGTCGGAGTGGCGCCGAACC |
| **Tmem119** | TTGGGGAGATGTTTCCTGGGTCCCTACACGACGCTCTTCCGATCTTGGCCGTGGGCATTC<br>CTGCTGAACCGCTCTTCCGATGGCCGTGGGGTCATTAAGGGCGCTGGAAC |
| **Cx3cr1** | AGGCAAAGACCACCAGGAGGTCCCTACACGACGCTCTTCCGATCTGCGGGCCGGTCATTC<br>CTGCTGAACCGCTCTTCCGAGCGGGCCGGTTGGTGTCCAGAAGAGGAAGA |
| **Gal** | TCTTCTCCTTTGCAGGCATCTCCCTACACGACGCTCTTCCGATCTGGGTCTGGGCCATTC<br>CTGCTGAACCGCTCTTCCGAGGGTCTGGGCGCTGTTCAGGGTCCAACCTC |
| **Olig1** | TTTTAATTGCCAGGGAGTGGTCCCTACACGACGCTCTTCCGATCTGCCCGCGGGGCATTC<br>CTGCTGAACCGCTCTTCCGAGCCCGCGGGGTGGAACACCCGCTTGGGTTA |
| **Gpx3** | TGCCATCTGGCCCCACCAGGTCCCTACACGACGCTCTTCCGATCTGCGCGGGTCGCATTC<br>CTGCTGAACCGCTCTTCCGAGCGCGGGTCGGTACCAGCGCATAACCGGTA |
| **Rasgef1b** | TGACACAGCCCTCGTTGAGGTCCCTACACGACGCTCTTCCGATCTGTGGGGCTTGCATTC<br>CTGCTGAACCGCTCTTCCGAGTGGGGCTTGATGGCCATTTGGAAGACGGT |
| **P2ry12** | GGCCCGGCTCCCAGTTTAGCTCCCTACACGACGCTCTTCCGATCTGGGCTGCGTGCATTC<br>CTGCTGAACCGCTCTTCCGAGGGCTGCGTGGGCACACCAAGGTTCTCAGA |
| **Apoc1** | TGTTCCGGACAAATCCGGGGTCCCTACACGACGCTCTTCCGATCTCGGGTTGGCGCATTC<br>CTGCTGAACCGCTCTTCCGACGGGTTGGCGTTATCCGGTATGCTCTCCAA |
| **Negative_1_human_BMP4** | CGTTACCTCAAGGGAGTGGGTCCCTACACGACGCTCTTCCGATCTGCGGCGTTTGCATTC<br>CTGCTGAACCGCTCTTCCGAGCGGCGTTTGGCATGTCAGGATTAGCCGAT |
| **Negative_2_human_Sox2** | CGTGAGCGCCCTGCAGTACATCCCTCACATCATAGAGGAATCGAGGTTGTGCGGCCATTC<br>AGTAGTCCAATAGAGGAATCGTTGTGCGGCCAGCCCATGCACCGCTACGA |

**Extra Table 2. The full list of the 10-base barcodes designed by hybrid block coding (n = 369)**

| | | | | |
|---|---|---|---|---|
| GCTCGGCTGG | CTGGGGGGCC | GGTTGGTTGC | GCTGCGGGCC | CGCTGGGTTG |
| GTGTCGCGGG | CCGGGGCGCG | GGCTTTCGGG | GGTTGTCCGG | CCGCGTGCGG |
| GTTGGCGTCG | TGGGCGCGTT | CGTCGCGGTG | CCGGGGTCGT | CCCGCGGGGG |
| GCTGGTGCGC | GTCGCCCGGG | CGGGTCCGTG | GGTGTTTCGG | GGGCGTTCTG |
| GGCGTGTGCG | GGCGGCCTGT | GCGGTCCTGG | GCTGGGTGTT | TCCGTGGGCG |
| TGGCCGGCGT | CGTTTGCGGG | GGTCGTCGGT | TGCGTCGGTG | GGCCTGGCCG |
| CGGCGGTTGG | TGGGGCTGCC | GTCCGGGTGT | GGTTTCGCGG | TTGGGCGTGC |
| TGCGGGCGGC | TGCCCGGGCG | TGGCCTGGTG | GTCTGGCTGG | GCGGCTCGTG |
| TTGCGGGGTG | TGCCGCGTGG | GGCGCTGGTG | TGGCCGTGGG | GTGGTGGCCG |
| CCTGGGGTGC | CGGGGCGTGT | CTGGCTGCGG | TGCGGGGCCT | CGGTGGTGTC |
| CGTGCCCGGG | TGGGGTCGTC | CCGCCGGGCG | CCTGGGGGCT | GCCGGGCCGC |
| CGTGCGGCTG | GGCCGCTCGG | GGGCGGCGCT | GGGTCTGGGC | GGCGCGCCTG |
| GGTCGGTCGT | GCGTCCGGGG | TTCGGGGGGT | GGGTGCCCTG | CTGGGGTTCG |
| TGGGCGTTCG | TGGGGGTTTC | GGGTTCGGTC | GCGGCTGTGT | TGGCTTGCGG |
| TGCTGTGGTG | GCGCCGGGTT | TGGTTCTGGG | CGGGGTCCCG | TGCGGCCGCG |
| GCTTGGGCGT | GCTCTGGGGT | GGTCGCCGCG | GCCCGCGGGG | GTCGGGTGTC |
| GTCGGTGTTG | GGGTCGGCTG | CGGCTGCCGG | GGCGTTGGCC | CTGGCGCGTG |
| GGCGCGGTGC | GGTTCTTGGG | TTGGGGTCTG | GCCGGCTGGC | GGTTCGCGTG |
| GGTTTGGGGC | CGCTGGGCGC | CGTGTGGGTC | CGGCCCGCGG | TGGCGGCCTG |
| GTTCGGGGCG | GGGTCCGTGT | CCGGGTGGGT | TGGGTGGCTC | TCGGGCTGTG |
| GGGCTCTGGT | TCGGGTGTGG | GGCTGGGTCC | GCGTGCGGTT | GTGCTGGTTG |
| TGCCTGGGGT | GGGCGTGCCC | GTGGTTGGTC | TCTCGGTGGG | TCGGCTTGGG |
| CGGCGCGGCT | GGGCTGGTGT | GGTCCGGGGG | CGGGGGCTTT | CGGGCCGTCG |
| TGGGGCCCGC | TGGTGTCGGG | GGCCGGTGTG | GCGCGGGTCG | GTGGTCGTGT |
| CCTGGCGCGG | CGGCGTTGCG | GCGGTGTGCT | CGCTTGGGCG | CGTCGGCGGC |
| CGCGGGTTGC | CTGTGGGCTG | TTGTGGGCGC | CGGGCCTGGC | GGGTGCTCGC |
| GGGTCGCTGC | GCCTGGTGGT | GTCTTGGGGT | TGTGGTGCTG | GGGCCTGGCT |
| CGCTCTGGGG | TTTGGGCGTG | GGTGGCGCCT | GTTGTCGGTG | GGGTGGTTCG |
| GGTGTGCTTG | GGCGCCTTGG | GCGGGTCTCG | CGGTGCCGGC | GCTCGTGGTG |
| GCTGGCTTGG | GCGCTTTGGG | GGCTCGTCGG | CGGGGTGCTT | CGTTGTGGCG |

| | | | | | | | | |
|---|---|---|---|---|---|---|---|---|
| GGTGGTTGCT | | GTGGGTTTGC | | TGTGCTGTGG | | CCGGGCGTTG | | GGCCCCGGGC |
| CGTGGGTCCG | | GCCGGTCGGG | | TGTCTCGGGG | | GTGGGGCTTG | | GTTGTGGGCT |
| GGTGTGGCCC | | GTGGGGCCGT | | CGGCGGGCCG | | CGGTGGCCGT | | CGGCCGCGGT |
| TGTGGCGGGT | | GGGCCTCCGG | | CGCGGCGGCC | | GTCGGGGCTT | | GCGTGGCGTG |
| CCGGTTCGGG | | GGTCGGGGTC | | CGGGCGGCGC | | GGGTGGTCTT | | TGGGGTTCGT |
| GCGCTCGGTG | | TGGGCCCTGG | | CGGTTGGTGC | | GTGGCCGGCC | | TTGGGCGGTT |
| GGCTGTGGGT | | CGGGTGTGGG | | CGCGGCTGGT | | CGGGTTGGCG | | GCCGCCGGGT |
| CTGGGGCTGC | | TGGGTTGGGC | | GCTTGGGTTG | | GTGCGCCGGC | | GCTGTGTGGG |
| GCCTCGGGTG | | GGGCGCTGTC | | TGTGGGCCGT | | TGCGGTGCGC | | GGTGCCGCGC |
| TCGTCGGGGT | | TGTGTGTGGC | | TGTCGGGTTG | | TTGCCCGGGG | | TCCCGGGCGG |
| GGCTGCTGCG | | TTGGGGTGGC | | GTTGCGTGCG | | GGGTTGTGCC | | GGCGCTTGGT |
| GGTGCTCGGC | | CTGTGCGTGG | | GGGCGTTTGT | | GCCGGTGGTC | | TGGGTCGGCT |
| GTGTGTGGCG | | GGCGGGCTCG | | TTGGTGCGCG | | CGTGTGGTGG | | CTTGGGGTTG |
| GGGCCGCTCG | | GGCCCGGTTG | | TGCTCGGGGC | | TGTGCGCGCG | | GTCGGTGGCT |
| TGGTGCGCGT | | CGCGGTGTGG | | GTGGCCGTTG | | GGGTTTGCGT | | CCGTGGGTGT |
| CGCCGGGGGG | | TCTGGGGTCG | | CGGCCGGGTC | | CGCGGCCCGG | | TGCTGGGTGT |
| GGCCTGTGGC | | GTGCTGTGCG | | GCCGTGGTGT | | GTGGCTCTGG | | TGGTGTGTGC |
| TGCGGGTCGG | | GTCGTGGCGC | | CCGGTGGCTG | | CTGTGTGGGC | | CTCGGGGCCG |
| TTGGCGCCGG | | GGGCGCGCTT | | GCGGCGCCCG | | GGTGGTCGTG | | GGCGTCGTCG |
| TTGTCGGTGG | | GGGTCCCGCG | | GCGGGCCGGT | | GTGGTCTGGG | | GCTTGGTGCG |
| TGTTGGGCGG | | GTGTGGGGTC | | GGTCTTGTGG | | CGGTGTTCGG | | TGGGGTGTCT |
| TGGGGCGCCG | | CTTGGTTGGG | | CGTTGGTGGT | | TCGGGTGGCC | | CTTGGGGCGT |
| GTCGCTGGGC | | GTGCGCTTGG | | GTGCGGTCGC | | GGCCGGCTGC | | GGTGCGTTGT |
| CTCGTTGGGG | | GTGTGCTGTG | | TGGTGGTGCT | | GTCGTGTTGG | | GTGGGTCGCC |
| GCGCGGGGGC | | GTGTTGTCGG | | GGGCTGCGTG | | TTGGGCCGGG | | CCCGGTGGCG |
| GGCCGTGGCG | | GCGGGTGCTG | | GGGTGTCGTT | | CGGGCCGGTT | | TCGGTTGGTG |
| CGTGCGGGGT | | TCGTGGGGCG | | GCTGCGCGGT | | TGTGGGGGCC | | TGCCGTTGGG |
| CGGGGTTTTG | | TCTTTGGGGG | | TTGGCTGGCG | | GCTTGCCGGG | | GTCCCGTGGG |
| GCGTTGGTGG | | CTGGTGGGGT | | GTGTCGGCGT | | TGCGCGTGTG | | GGCGGGTTTT |
| GCGGGCGGCG | | GTTGGCGGGC | | TGTCGTGGGC | | GGCGGGTCCC | | GTGGCGGCTC |
| GGCGTGCGGT | | GTCGGTTCGG | | TGGCTGGTCG | | GCGCCGGCGG | | CGGCTCGGGC |
| TTCTGGTGGG | | GCGGTCGCGC | | GCGTGTTTGG | | GCCTGGGCCG | | GGGTCGTGGT |

| | | | | | | | | |
|---|---|---|---|---|---|---|---|---|
| GGTGCGTGTC | | GTGGCGTGTT | | CTGGGTGGTG | | TTGCTGGGGC | | TTTGCGGGGG |
| CGGGTGCGCC | | GTCGTGCGTG | | TGGGTGCTGT | | GTGTGGCCCG | | GGGCTTCGGC |
| CTCGGGCGGG | | GTTCTGGCGG | | CGCGGGCGCT | | GGTCGGGTCT | | GCCGTTGCGG |
| GCCGGGTCTG | | CTTTGGGGGG | | TCGTGGCTGG | | CGGTCGGGCT | | TCGCTGCGGG |
| TCGCGCGGGT | | GGCGTCGCGT | | CGCGCGCTGG | | CCGCGCCGGG | | GGCGGCGCTC |
| TGTGTTCGGG | | GCGGGGGCCT | | CCGCGGTGTG | | GCGGTGTTGC | | GCGTGCGTGC |
| GTGCGTCGTG | | GCCCGGCGCG | | GGTGCGGTCG | | GGCGCGGGCT | | TGGTGCGTTG |
| GCTGCTGGGG | | GTGGCGGTCT | | CCGTCGTGGG | | GGTGGCTCTG | | GTGCCGGTGC |
| GTGGGCGCGG | | GCGGGGGTTC | | TCGGTGTCGG | | TTGTGGCGGT | | CTGCGGTGGT |
| CCGGCGCGGC | | GTGGGCTGCT | | GCGTTGCGGC | | GGTCGCGTGC | | GCGGCGTTTG |
| GGGTTTGTCG | | TCGGCCGCGG | | GGTGGTGTTC | | GGTGGGCCTC | | GTTGTGCGGC |
| CGGGTGGCCT | | GTGCTTGGGT | | TCGGCGGGTC | | GGTTGGCGCC | | |

### Extra Supplement PDF (following pages)

Transcript density of SPRINTseq in comparison with FISH images from Allen Brain Atlas.

Transcript density in comparison with ABA FISH images.

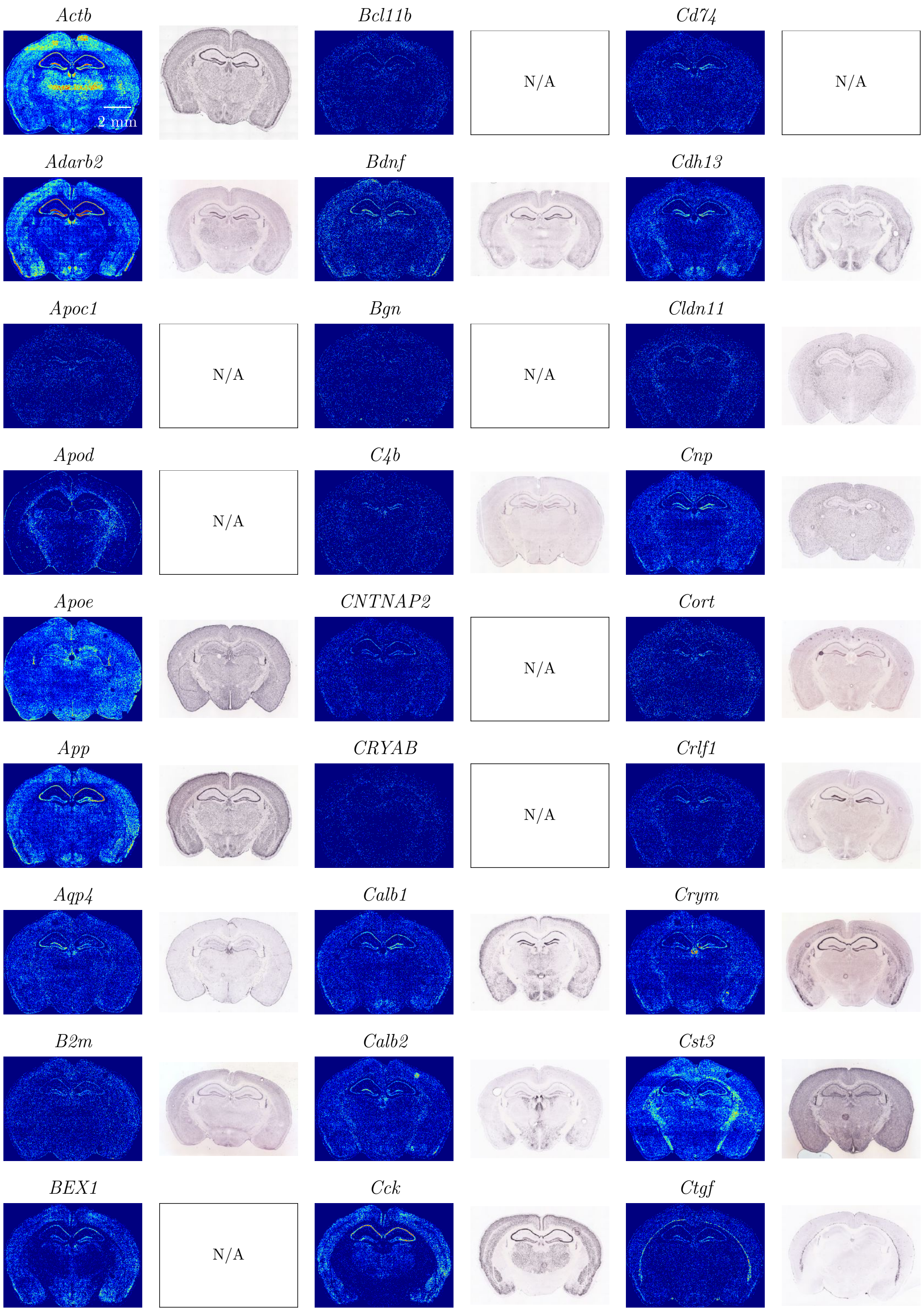

Continued on the next page

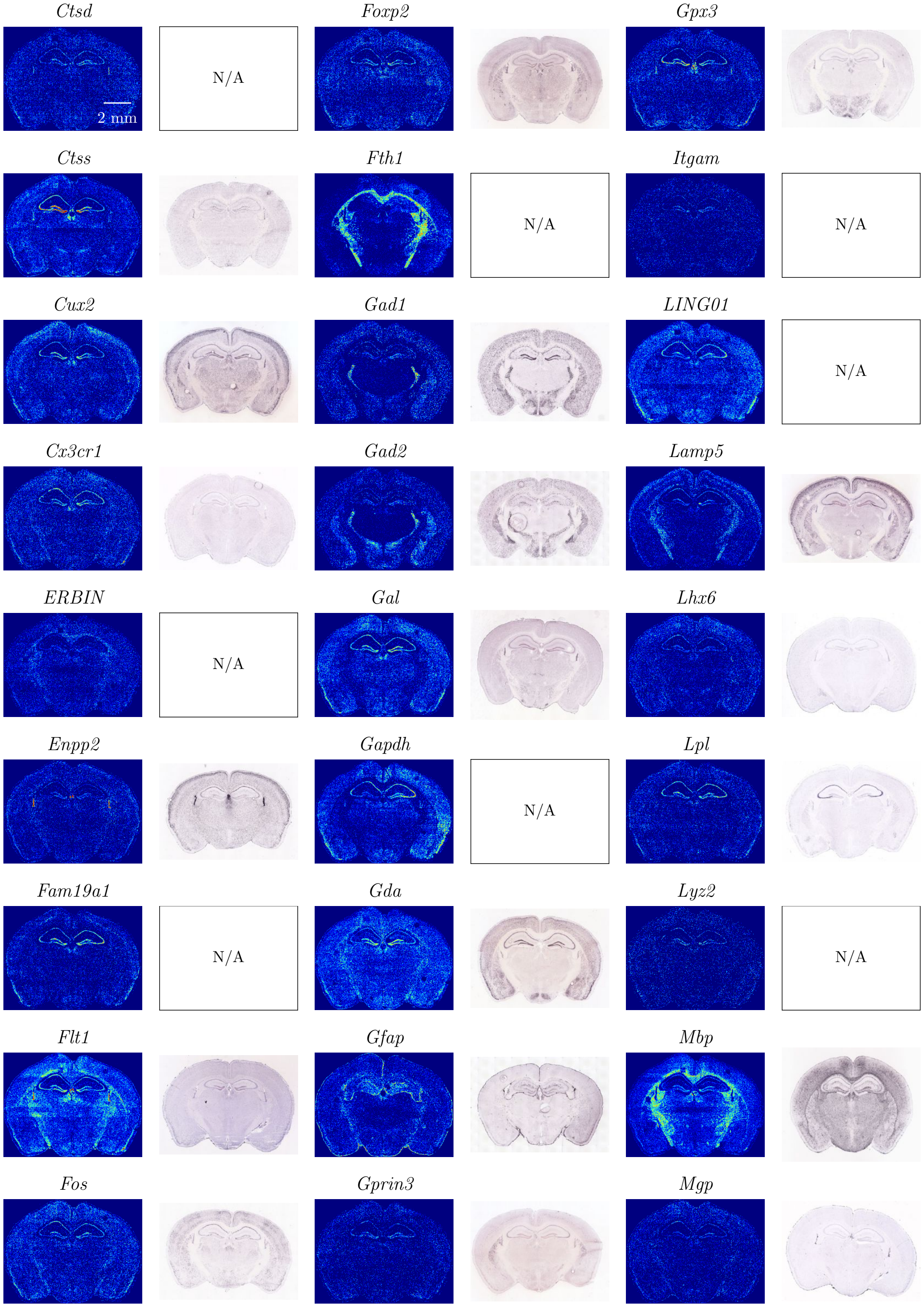


Continued on the next page

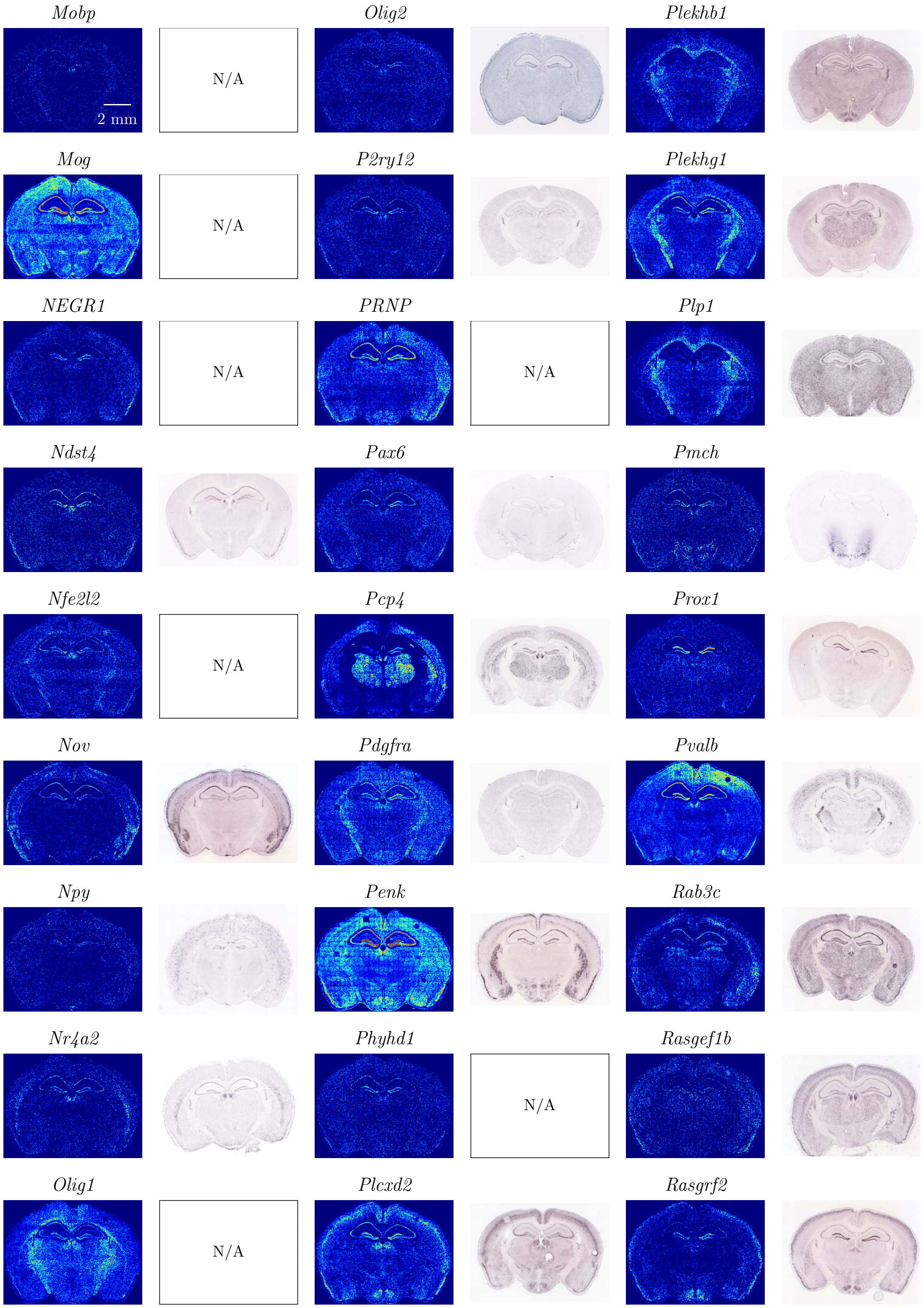


Continued on the next page

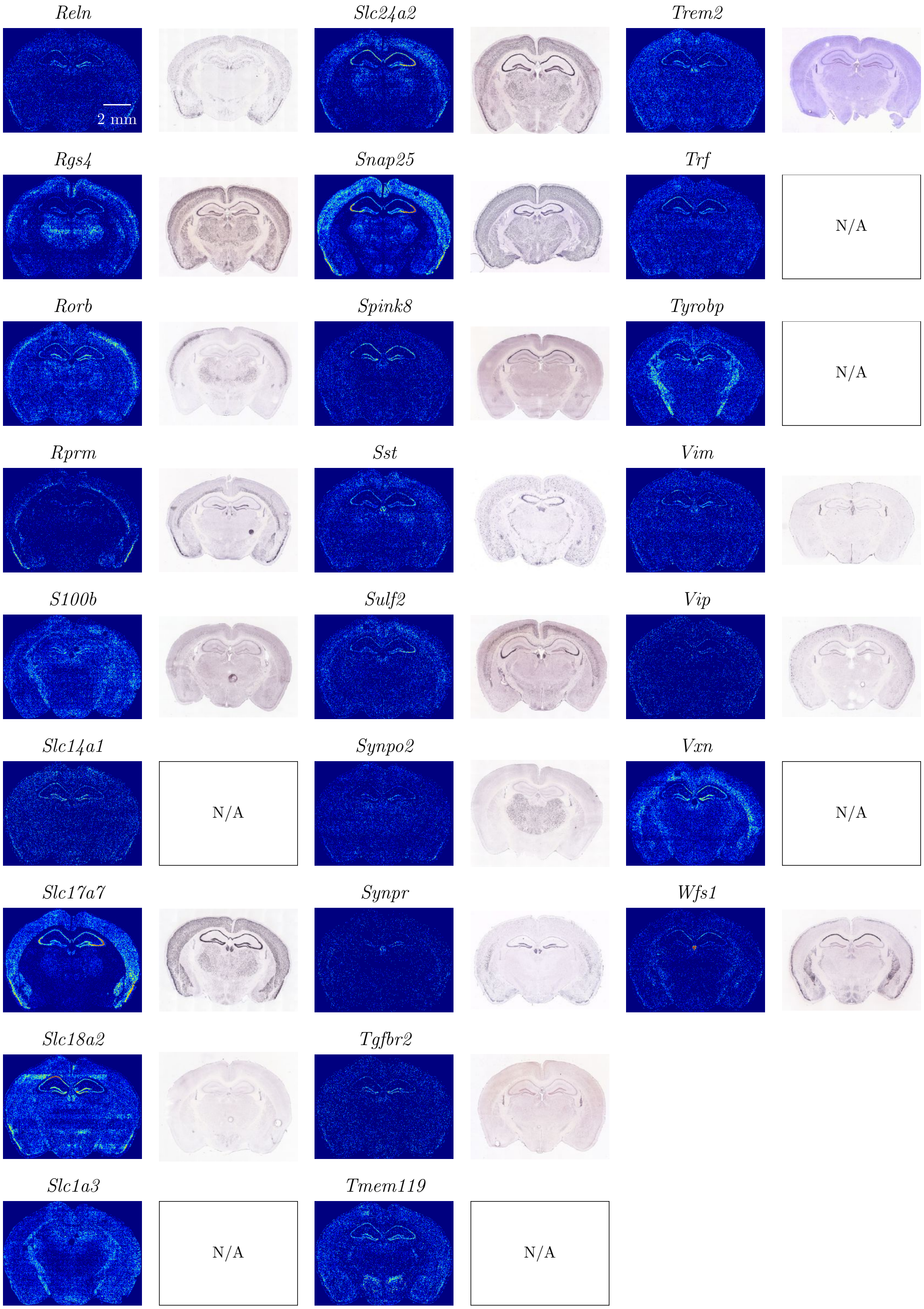
Reln
2 mm
Slc24a2
Trem2
Rgs4
Snap25
Trf
N/A
Rorb
Spink8
Tyrobp
N/A
Rprm
Sst
Vim
S100b
Sulf2
Vip
Slc14a1
N/A
Synpo2
Vxn
N/A
Slc17a7
Synpr
Wfs1
Slc18a2
Tgfbr2
Slc1a3
N/A
Tmem119
N/A
